# CA1 ensemble plasticity is coupled to context change and modulated by task familiarity

**DOI:** 10.1101/2024.08.19.608588

**Authors:** Branislav Krajcovic, Daniela Cernotova, Ales Stuchlik, Stepan Kubik, Jan Svoboda

**Author notes:** Corresponding author: Branislav Krajcovic, Department of Physiology, Second Faculty of Medicine, Charles University, Plzenska 311, 150 06, Prague 5 Motol, Czechia. contributed equally (Stepan Kubik, Jan Svoboda).

## Abstract

When should plasticity mechanisms get recruited (stability-plasticity dilemma)? Environments change over time, re/entering a given context increases uncertainty and predicts the need for updating. Hippocampus (HPC) is key to tracking context change but also navigation in relation to moving targets. CA1 ensembles expressing immediate-early genes (IEGs) are contextually specific, while the amount of IEG expression correlates with HPC-dependent task demands. However, task effects on the IEG-expressing ensembles *per se* remain unclear. In three experiments, we tested the effect of context change and HPC task demands on CA1 IEG+ ensembles in rats. Experiment 1 showed that the IEG+ (*Arc*, *Homer1a* RNA) ensemble size drops to baseline level during uninterrupted 30 min exploration, reflecting familiarization and decreasing uncertainty, unless context change is present; the ensemble sizes reflect both context identity and context change. Experiment 2 showed no evidence of task-specificity of IEG+ ensembles during highly HPC-dependent mobile robot avoidance nor HPC-independent stationary robot avoidance. Experiment 3 replicated the findings of Experiment 2 for c-Fos protein. Nonetheless, the data suggest that ensembles shrink with task mastery/familiarity and grow with novelty presented by acquisition of behavioral extinction. Overall, our results shed light on the temporal dynamics, and the context and task control of CA1 IEG+ ensembles. The present results and the relevant literature suggest that context change resets the ensemble of IEG-expressing CA1 neurons and novelty delays the time-dependent ensemble shrinking.

**HIGHLIGHTS:**

- Plasticity and learning rate should reflect novelty and familiarity, i.e. uncertainty
- Change of context and task requirements increase uncertainty
- Familiarization with context and task reduces uncertainty
- For context, this pattern is matched by dynamics of IEG+ ensembles in CA1
- Task demands have modulating influence on CA1 IEG+ ensembles

**GRAPHICAL ABSTRACT:** 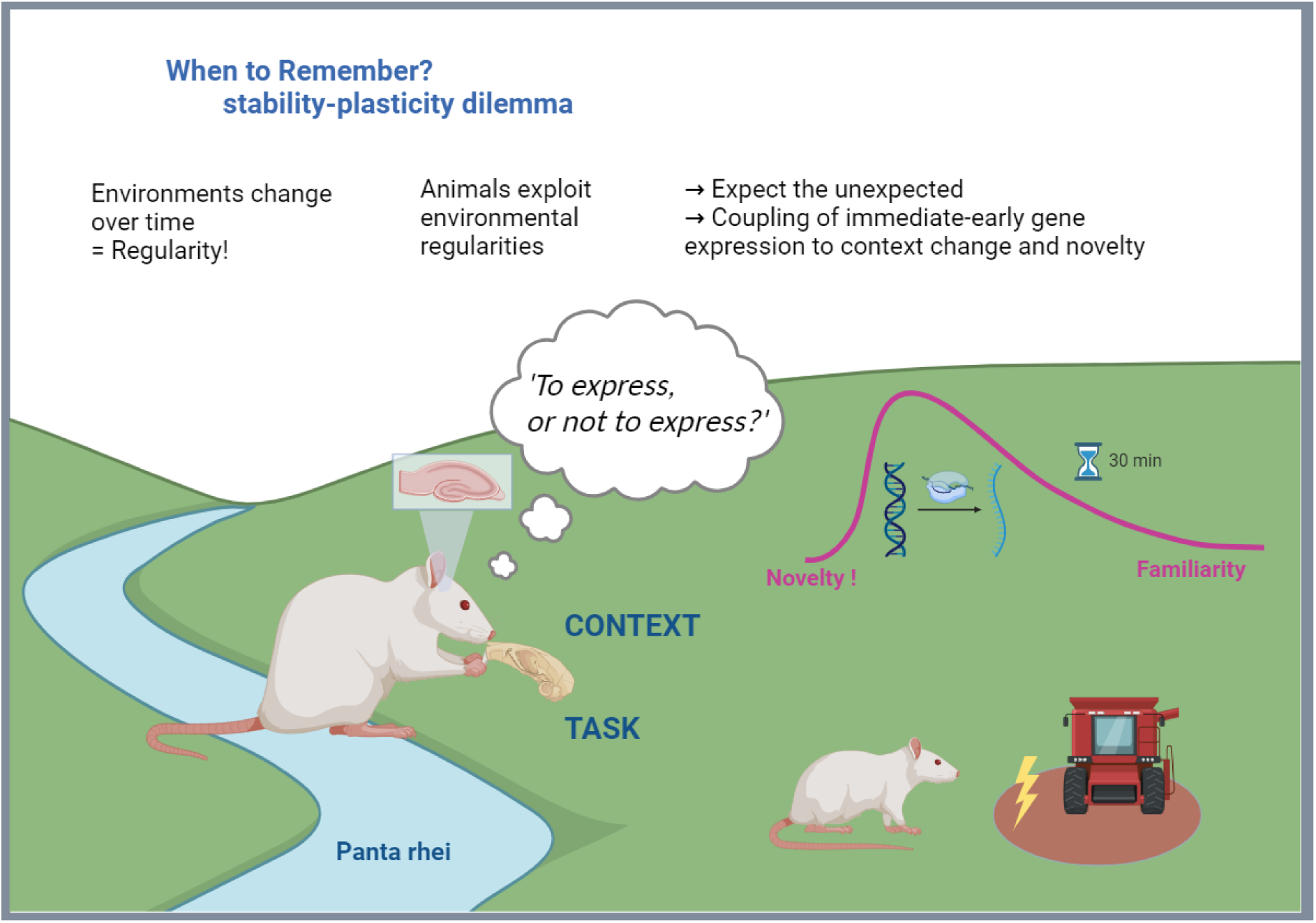

## 1. INTRODUCTION

‘Everything flows’ as Heraclitus noted. For behavior to remain adaptive in a changing world, learning is essential since it allows updating of knowledge and keeping predictions accurate. However, plasticity mechanisms enabling learning and memory cannot work constantly at their maximum capacity as this would result in saturation rendering plasticity fruitless and compromising memory (1,2). Consequently, neural systems face the stability-plasticity dilemma (3): when should the plastic processes be recruited? The naive, but not untrue, answer is that ‘when something new and behaviorally relevant happens or is about to happen’. The problem then becomes one of uncertainty estimation and learning rate adjustment(4). Although changing, the environment is not completely random. Organisms evolved embedded in and constantly interacting with their surroundings (5). Biological systems are coupled to environmental regularities and exploit invariant or statistically regular relationships to their advantage (6). One such regularity that might aid the stability-plasticity dilemma is the relationship between continuous presence in a given environment and decrease of novelty. Generally, the longer we continuously stay in the same environment, the more familiar we get with its current state, the uncertainty decreases, and so does the need for updating. Conversely, the very fact that environments change over time means that re/entering an environment increases uncertainty. Neural systems can use the change between contexts, and the associated novelty, as an indicator of uncertainty to trigger plasticity mechanisms that enable knowledge updating and improve inference (4).

The hippocampus is crucial in processing contextual information (7–10), tracking the continuity and change of context (11). Change of spatial context rapidly and transiently triggers expression of immediate-early genes (IEGs) (12–15). The IEG expression is the first line of genetic response to neuronal activity, enabling lasting synaptic adjustments and long-term memory: the IEGs *Arc* and *Homer1a* give rise to effector proteins of the postsynaptic density, whereas the IEG *c-fos* acts as a transcription factor (16,17). The hippocampal CA1 ensembles expressing *Arc* and *Homer1a* are specific to spatial context (18,19). Although IEG expression levels correlate with task demands, learning, and performance (12,20,21), the specificity of CA1 IEG+ ensembles *per se* to task demands or performance is less clear (22). In three experiments, we investigated the dynamics of IEG-expressing ensembles in hippocampal CA1 as it relates to novelty, familiarity and re-visiting of spatial context, but also the effect of unexpected task demands in a familiar environment on CA1 ensemble size and stability.

In Experiment 1 (Exp1), we investigated the dynamics of *Arc+* and *Homer1a+* CA1 ensembles over a 30-minute period with respect to timed introduction to novel or familiar context, or continuous presence in a given environment. In Experiment 2 (Exp2), we examined how unexpected task demands in a familiar environment affect IEG expression on the ensemble level. We used the ‘robot avoidance task’, which allows adjusting the hippocampal load: avoidance of a mobile, but not stationary, aversive robot is seriously impaired by transient hippocampal inactivation (23). While the primary RNA transcripts used in the first two experiments provide the most immediate view of plasticity-related activity, labeling IEG protein captures a longer time window. In Experiment 3 (Exp3), separate groups of animals received the same treatment as in Exp2, but we examined c-Fos protein for potential differences on a longer timescale.

## 2. METHODS

### 2.1. Animals

A total of 118 adult male Long-Evans rats bred in the Animal Facility of the Institute of Physiology CAS were used for all experiments. All animal procedures were approved by a local ethical committee and complied with the Animal Protection Code of the Czech Republic and with the EU directive on the use of laboratory animals (2010/63/EC).

In Experiment 1, twenty-four rats were used. Rats were maintained at standard conditions (22°C, 50–60% humidity, 12 h light/dark cycle, food and water ad-libitum), housed by 2-3 in transparent cages, and tested at 4-5 months of age. The rats received 3 handling sessions prior to the test session, but no habituation, so the testing environment(s) were novel at the time of the test for behavioral induction of IEG expression. After the last handling session, the day before the test, rats were housed individually in opaque cages to minimize confounding IEG expression due to social interaction.

In Exps 2 and 3, forty-eight and forty-six rats, respectively, received the same behavioral treatment. Rats were housed by two at the same standard conditions as in Exp 1 and tested at 2–3 months of age. During these experiments, rats were food-restricted and kept at 85-90% of their pre-restriction weight to motivate exploration and prevent passive coping. Three days before the robot avoidance training, a fine hypodermic needle (diameter 0.6 mm, 30 mm long) was implanted through a skin fold on the back of the neck, then blunted and twisted into a loop on its endpoint to prevent slipping out. The needle implant allowed footshock delivery during training by closing a constant-current circuit with the grounded arena surface. Shock intensity was set individually between 0.2-0.6 mA to motivate escape response but avoid freezing behavior.

Overall, 8 animals were excluded from experiments 1 and 3, resulting in a total of 110 animals included in the analyses for all the experiments reported here. In Exp1, one cage control animal was excluded due to exceptionally many *Arc+* neurons (36%). Such high expression in a caged control animal indicates that the rat underwent uncontrolled strong disturbance shortly before the sacrifice. Both the literature (18,19,24) and our own data (15,25) repeatedly and reliably demonstrated that the baseline expression of *Arc* involves only around 5% of the dorsal CA1 neurons in quiet conditions; we therefore excluded this animal from all statistical analyses. In Exp3, seven animals were excluded from the analyses: 1 rat was due to loss of data, and 6 rats due to sample damage during tissue processing. Additionally, in the exploratory analysis of behavioral extinction in Exp3, two animals from the MS group were not included due to missing data on the number of virtual shocks on test day 5; however, these two animals were included in the main analysis as data on shocks for training days 1-4 were intact and the number of virtual shocks on day 5 was not part of the main analysis of c-Fos expression.

### 2.2. Experimental design

#### 2.2.1. Experiment 1: Behavioral induction of context-specific *Arc* and *Homer1a* expression: effect of continuous exposure compared to repeated sessions

##### 2.2.1.1. Apparatus

The test session(s) for IEG induction in Experiment 1 were carried out on an elevated circular metallic platform (diameter 82 cm) equipped with 3 identical objects (brown plastic cylinders) enclosed with a transparent plexiglass wall (height 50 cm), located in a quiet room under constant low lighting (220– 250 lux) – context A (**Fig. 1A**). Some of the animals also experienced context B (**Fig. 1B**), a square arena (side 70 cm) with 40 cm high white walls, equipped with 3 identical wooden blocks, and located in a different experimental room; the lighting conditions were again kept low. To promote the distinctiveness of contexts A and B, the apparatuses were cleaned using 70% ethanol and soap solution, respectively.

**Figure 1.**
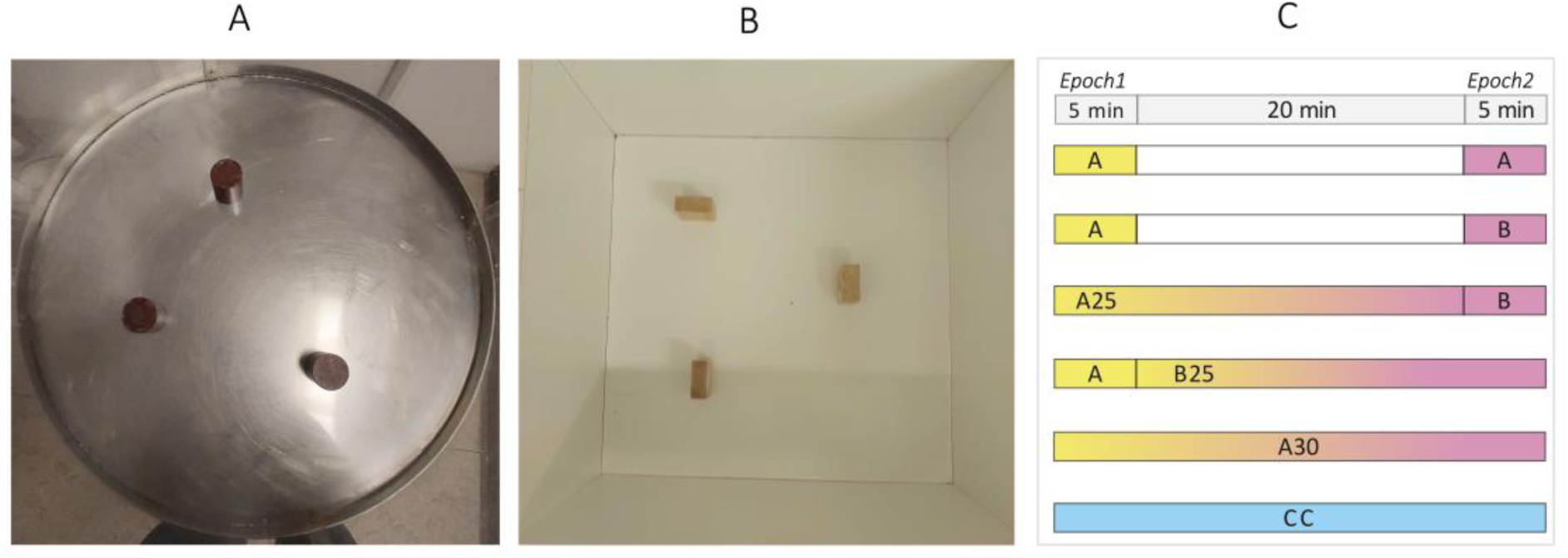
Behavioral design and experimental apparatuses used in Experiment 1. **A.** Context A was a circular metallic arena (82 cm diameter) with a 50 cm high transparent plexiglass wall and three identical plastic columns. **B.** Context B was a square white arena (70cm side) with 40 cm high walls and three wooden blocks. **C.** Twenty-four rats were split into 6 groups with different behavioral treatments. A/A rats were exposed to context A twice for 5 min with a 20-minute interval in the home cage. A/B rats were exposed to different contexts A and B for 5 min each and a 20 min interval in the home cage. A25/B rats were exposed to context A for 25 min and then immediately transferred to context B. A/B25 rats were exposed to context A for 5 min and then immediately transferred to context B for 25 min. A30 rats were exposed to context A for a continuous 30 min. Home cage control (CC) rats remained undisturbed in their home cages. Neuronal activity in the first 5 min of the test (Epoch1) was tagged by the *Homer1a* signal, and activity in the last 5 min (Epoch2) is tagged by the *Arc* signal.

##### 2.2.1.2. Behavioral design

On the testing day, rats were quietly transferred to a holding room and assigned to one of the following 6 groups based on which of two distinct novel contexts A and B, and for how long they visited: A/A, A30, A/B25, A25/B, A/B, and CC (**Fig. 1C**). Locomotor activity was analyzed automatically with EthoVision XT 17 (Noldus, Netherlands).

#### 2.2.2. *Arc/Homer1a* catFISH

*Arc/Homer1a* catFISH enables comparing intranuclear pre-mRNA IEG signals induced by behavior during two distinct time epochs. Expression of both IEGs is triggered at the same time, but due to different lengths of the transcripts and target location of the antisense probe (described in (24)) the catFISH allows mapping plasticity-related ensemble activity during the first (Epoch1, *Homer1a*) and last (Epoch2, *Arc*) ∼5 minutes of a given 30-min period.

##### 2.2.2.1. Tissue Processing and Labeling

Immediately after the test session, rats were deeply anesthetized with isoflurane, brains were quickly (<2 min for Exp1, 2.5 min for Exp2) removed and flash-frozen in an isopentane bath equilibrated in dry ice/ethanol slurry. Cryosectioning (20 μm) and double label (*Homer1a*, *Arc* RNA probes) fluorescence *in situ* hybridization for dorsal CA1 was performed (**Suppl. 1**) as described previously (14,19,25). Eight hemispheres from multiple groups were arranged into a block for cryosectioning to maximize between-group comparisons on each slide.

##### 2.2.2.2. Image Acquisition and Sampling

In each left dorsal CA1, three microscope fields of view (FOV) were selected, starting at the apex of dorsal CA1 and following in the lateral direction (proximal CA1). For each FOV, a confocal Z-stack of 20 images at 1 μm z-step size was acquired. Details on microscopy protocol are in the supplements (**Suppl. 2**) Images were analyzed semi-automatically using a custom macro for ImageJ as described previously in Kubik et al. (14). Regions of interest (ROIs) for neuronal nuclei were selected only in the middle 20% of each Z-stack to avoid IEG signal cut-off. Both image acquisition and subsequent analysis were performed by a blinded experimenter. Two slides per rat provided six CA1 Z-stacks per rat resulting in a total of 144 Z-stacks. No image exclusions were made from this dataset. A total of 9631 nuclei were counted with an average of 401.3 nuclei per animal in Exp1.

##### 2.2.2.3. Similarity Score

Similarity scores were calculated according to Vazdarjanova and Guzowski (19). They express the overlap of IEG+ expressing ensembles active during the two epochs normalized to the cell population size; a value of 0 indicates random overlap of statistically independent ensembles, while a score of 1 indicates 100% perfect non-random overlap i.e. exactly the same ensemble active in both epochs. Similarity score calculation: ss = diff(E1,E2)/(least(E1,E2) - random overlap(E1,E2)) = (double+ - E1*E2)/(least(E1,E2) - E1*E2) where diff(E1,E2) = difference between observed (double+) and expected overlap (E1*E2); E1 = *Homer1a*+ % (Epoch1); E2 = *Arc*+ % (Epoch 2); least(E1,E2) = smaller of E1, E2%; double+ = *Arc*&*Homer1a*+ %.

#### 2.2.3. Experiment 2: *Homer1a* and *Arc* RNA expression and avoidance of a stationary or moving robot

##### 2.2.3.1. Apparatus

Behavioral parts of Exp2 and 3 (**Fig. 2**) were performed in a quiet room under constant dim lighting (220–250 lux) on an elevated circular metallic arena (130 cm diameter) enclosed with a transparent plexiglass wall (50 cm height). The robot was custom-made, three-wheeled, car-like in shape (length 17 cm, width 12 cm, height 10 cm). The robot was programmed (Arduino, Italy) to move straight until it bumped into a wall, pause for 15 s, move 10 cm backward, turn at a random angle between 100 and 200 degrees, and continue onwards. The robot’s speed was set to either 4 cm/s (moving) or 0 cm/s (when stationary). Custom-made software (Kachna tracker, author Tomáš Mládek) tracked the position of both the rat wearing a harness with 2 colored LED markers, and the robot. The starting position of the robot changed for each session; rats were placed in the arena as far as possible from the robot to allow avoidance. A motorized overhead feeder scattered pasta rice over the arena surface during all training sessions to promote exploration.

**Figure 2.**
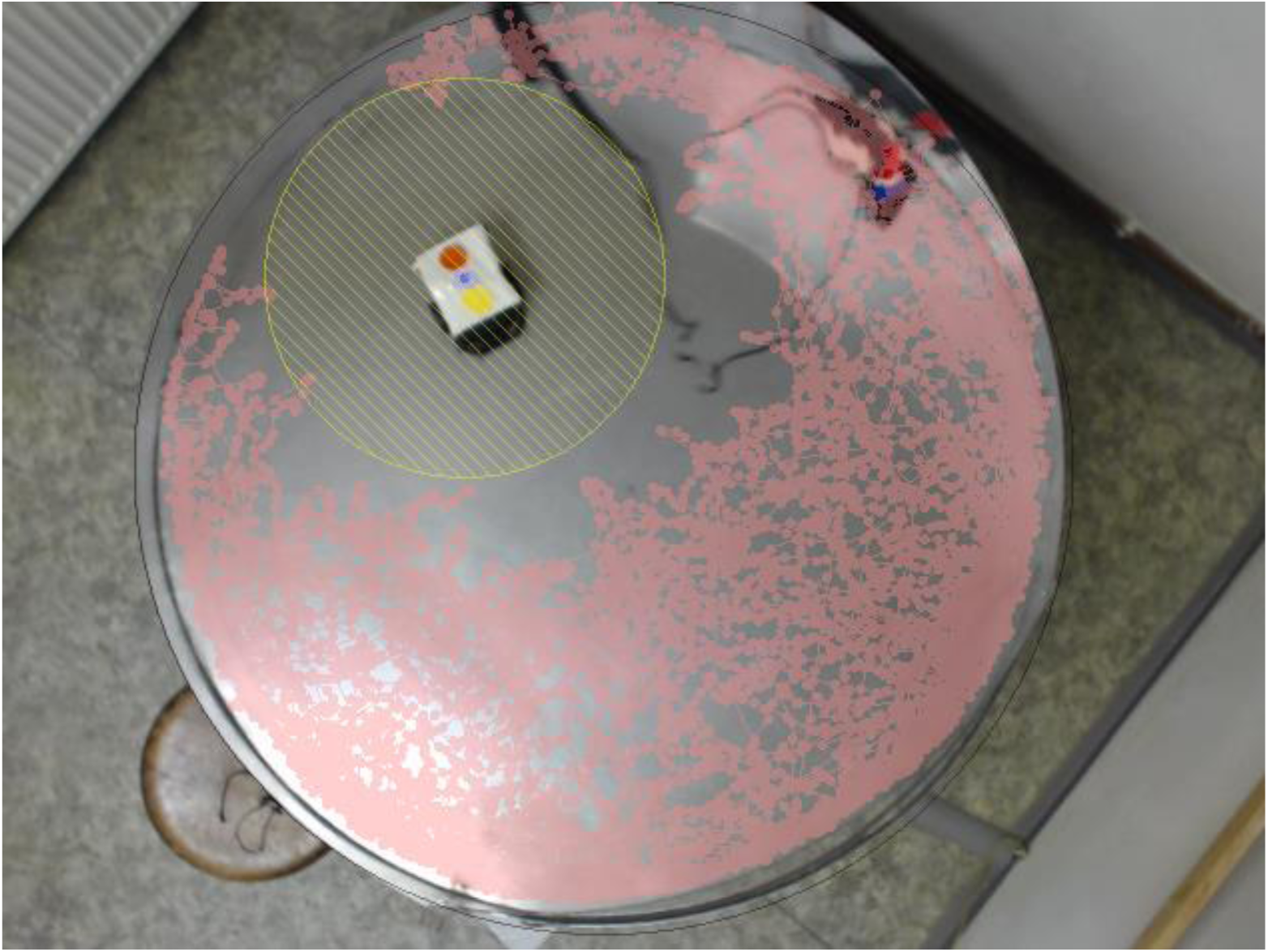
Robot avoidance in Experiments 2 and 3. Overhead view of the experimental arena and a rat avoiding a stationary robot at the end of a 20-minute session. Yellow hatchings mark the shock zone around the robot, the rat’s trajectory is shown in pink.

##### 2.2.3.2. Behavioral design

In both Exp2 and Exp3, rats were handled for 2 days and, on each day, also habituated to the arena for 5 minutes (no robot present, no shock delivered, only foraging for pasta rice).

All the groups with their respective labels are described in **Tab. 1**. The experimental rats learned to avoid a stationary (S) or moving (M) robot (20 min/day for four days). On day 5, they were tested in either the same (SS, MM) or the opposite condition (SM, MS) with the shocks turned off. In addition, ‘active controls’ received four days of pseudo-training with a stationary robot but no shocks (nsSS) to account for the effects of transfer and exploration without explicit learning requirements. ‘Passive controls’ remained in their home cages during the test: some of these rats previously received training with either moving robot (MC) or stationary robot (SC), or no training at all (CC); all these rats unanimously showed low IEG expression, and they were pooled and treated as one CC group.

**Table 1.**
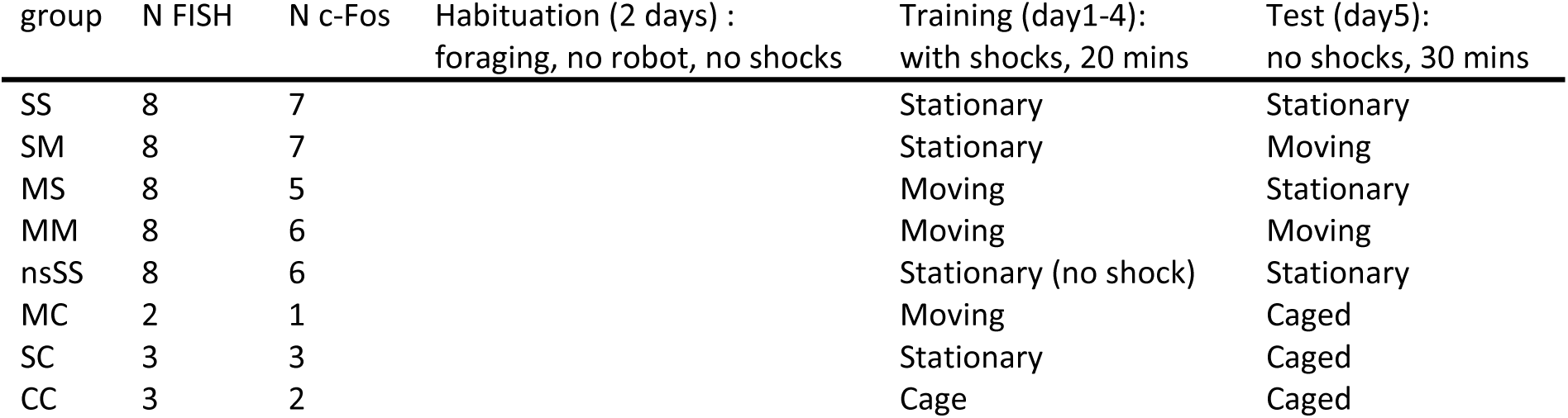
Design of Experiments 2 and 3. Treatment groups are shown in rows. The first column provides group labels based on the robot’s condition in training and testing (e.g. SM=robot Stationary during training, Moving during test). Columns 2 and 3 show the number of animals in each group in Exp2 and 3, respectively. Behavioral treatment: 2 days of habituation (col. 4); 4 days of training with the robot (col. 5), test on 5th day for IEG assessment (col. 6). Descriptions in columns 5 and 6 refer to the robot’s condition during that phase (training or testing). The nsSS group served as an ‘active’ control: rats explored the arena with a stationary robot but were not required to avoid it since shocks were turned off (virtual shocks recorded). Groups in the last three rows remained in their home cages during the testing period (cage) and were treated as a single ‘passive’ control group (CC).

| group | N FISH | N c-Fos | Habituation (2 days) :<br>foraging, no robot, no shocks | Training (day1-4):<br>with shocks, 20 mins | Test (day5):<br>no shocks, 30 mins |
| --- | --- | --- | --- | --- | --- |
| SS | 8 | 7 |  | Stationary | Stationary |
| SM | 8 | 7 |  | Stationary | Moving |
| MS | 8 | 5 |  | Moving | Stationary |
| MM | 8 | 6 |  | Moving | Moving |
| nsSS | 8 | 6 |  | Stationary (no shock) | Stationary |
| MC | 2 | 1 |  | Moving | Caged |
| SC | 3 | 3 |  | Stationary | Caged |
| CC | 3 | 2 |  | Cage | Caged |

During training, the rats received a mild electric footshock (0.2-0.6 mA) whenever they got closer than 25 cm to the robot. After the last training session, rats were housed individually in opaque cages, and stayed in a separate room overnight until the test. The test lasted 30 minutes with the robot present, but no shocks were delivered (virtual shocks recorded) to prevent confounding IEG expression due to the shock itself (26).

##### 2.2.3.3. *Arc/Homer1a* catFISH image data sampling

The Arc/Homer1a catFISH procedure was identical to that described for Exp1, only the image data sampling was slightly different. In Exp2, four brain sample sections per animal were used. In total, 576 Z-stacks were acquired, of which 35 Z-stacks were excluded from image analysis due to tissue damage or weak staining of a sample slide. Overall, an average of 11.3 Z-stacks were analyzed per animal, and on average, 0.7 Z-stack per animal was excluded from image analysis. Due to group randomization on each sample slide, the excluded Z-stacks were evenly distributed across groups. In total, 21484 nuclei were analyzed, with an average of 448 nuclei per animal. Details on cell counts are in **Suppl. 4**.

#### 2.2.4. Experiment 3: c-Fos protein expression and the robot avoidance

##### 2.2.4.1. Behavioral design

The c-Fos protein expression was assessed in a separate cohort of animals due to different tissue processing requirements; experimental design and groups exactly matched the design of Exp2. After the testing session, rats were returned to opaque cages for 50 minutes to allow peak accumulation of c-Fos protein.

##### 2.2.4.2. Tissue Processing for c-Fos Expression Assessment

Fifty minutes after the session, rats were deeply anesthetized with isoflurane and transcardially perfused with 0.1 M PB and then 4% paraformaldehyde. Brains were post-fixed in PFA, dehydrated in 30% sucrose, quick-frozen, and stored at -80°C. Forty micrometers thick slices were cut (Leica cryostat). Every 6th cut was stained for DAB (**Suppl. 4**).

##### 2.2.4.3. c-Fos Image Acquisition and Analysis

In each dorsal left CA1, one FOV was selected, starting at the apex of dorsal CA1 and following in the lateral direction (proximal CA1). All images were acquired on an Arsenal microscope LP 3015 using a DinoEye C-Mount Camera. Images were analyzed using ‘interactive machine learning for (bio)image analysis’ software (ilastik; version 1.4.0b27) (27) by applying Pixel and Object classification workflow separately. All segmented cells were manually checked by a blinded experimenter. Manual corrections involved removal of artifacts, clear false positives and negatives. In total, 5–9 images were counted for each rat, depending on the amount of stained hippocampal slices. One image included approximately 1 mm of CA1 in length.

#### 2.2.5. Analysis of behavioral extinction during test session in Experiments 2 and 3

Since the shock delivery was turned off during the test session (day 5), some rats experienced behavioral extinction of the avoidance response. We performed an exploratory analysis, set extinction thresholds for Epoch1 and 2 of the test session, categorized the animals and assessed IEG expression. We used training day 4 data to calculate one-sided tolerance intervals (TI, 99% proportion of population, 95% confidence level) and used the TI upper-bound as thresholds on the number of virtual shocks received during Epoch1 and 2 of test day 5. Animals from Exp2 and 3 were pooled for TI estimation as the behavioral design was identical. The TI estimates the range of expected values within a specified proportion of a distribution with selected level of confidence: if the number of virtual shocks received on day 5 by a given animal exceeds the day 4 TI we can have 95% confidence that such extreme data would occur in only 1% of day 4 distribution. In such a case, extinction of the avoidance response can be assumed as present.

To set extinction thresholds for Epoch1 (test session t=0-5 min), we calculated one-sided tolerance intervals (99% population, 95% confidence) for the first five minutes of day 4. We separated rats into two groups: stationary robot training (SS+SM, N=30, threshold ‘S-5’) and moving robot training (MS+MM, N=27, threshold ‘M-5’). Thresholds S-5 and M-5 were applied to rats based on their robot exposure on day 5. For Epoch2 (test session t=25-30 min), we calculated one-sided tolerance intervals (99% population, 95% confidence) for the number of shocks received during the entire training session on day 4 (t=0-20 min). Groups were again split into stationary (SS+SM, N=30, threshold ‘S-20’) and moving robot (MS+MM, N=27, threshold ‘M-20’).

Next, we applied the thresholds to categorize the animals (Epoch1-Epoch2, ‘Av’ Avoidance, ‘Ext’ Extinction) as Av-Av, Av-Ext, Ext-Ext, Unclear. For Epoch1, animals receiving fewer virtual shocks than the S/M-5 threshold were labeled ‘Av’, and those exceeding it were labeled ‘Ext’. For Epoch2, rats not exceeding the S/M-5 threshold during the whole test were labeled ‘Av’, and those crossing the S/M-20 threshold at 20 minutes were labeled ‘Ext’. Rats not fulfilling these criteria were excluded from further IEG analysis (labeled ‘Unclear’) as the assessment of avoidance/extinction after the 20th-minute mark of the test could not be directly compared to day 4 data.

Although in Exp3 we evaluated c-Fos protein for the whole test session, and not individual epochs, we classified animals the same way as in Exp2 (Epoch1-Epoch2). We presumed differences in c-Fos expression might exist between animals, in which extinction occurred early or late in the session.

#### 2.2.6. Data analyses

Two-way ANOVA with repeated measures on the two behavioral epochs was used to analyze locomotion and IEG expression in Exp1 and 2. Two-way ANOVA with repeated measures on the four days of training was used to assess robot avoidance in Exp2 and 3. One-way ANOVA was used to analyze differences in similarity scores and c-Fos counts across the groups. Šídák’s and Holm-Šídák’s multiple comparison tests were used as appropriate. These analyses were performed using GraphPad Prism 10.0. To estimate the effect sizes, we used the DABEST package v2023.03.29 (28)[26] in Python v3.11 to calculate bias-corrected and accelerated (BCa) confidence intervals (CI) (29); 5000 bootstrap samples were taken. Custom Python code was used for some of the plots; minor graphical adjustments were done in Corel Vector and CorelDraw. Behavioral data were analyzed with CM_manager 0.5 software (30), DABEST package or custom Python scripts. For the analysis of IEG expression based on the extinction groups we used only descriptive statistics and did not proceed with statistical testing as these groups were created *ex post* from the already analyzed dataset.

## 3. RESULTS

### 3.1. Experiment 1 - Effect of context change or continuous presence on the size and overlap of IEG-expressing neuronal ensembles

An overview of data and effect size estimates for Exp1 is provided in **Tab. 2**.

**Table 2.** Effect Sizes for Experiment 1. Locomotion, Ensemble size and overlap.

| Group | Epoch1 | Epoch2 | Paired mean difference<br>Epoch2 minus Epoch1 (95% CI) |
| --- | --- | --- | --- |
| CC | x | x | x |
| A/A | 20.8±3.1 | 18.2±2.0 | -2.61 [-7.19, 0.19] |
| A/B | 20.3±4.0 | 20.2±2.3 | -0.11 [-4.47, 3.73] |
| A25/B | 21.0±1.5 | 20.0±1.1 | -0.96 [-1.31, -0.40] |
| A/25B | 19.4±1.1 | 1.5±1.6 | -17.9 [-18.6, -17.3] |
| A30 | 21.9±3.0 | 5.4±3.7 | -16.5 [-17.8, -15.6] |

***Ensemble size: Proportion IEG+ cells (mean, std)***
| Group | *Homer1a+*<br>(Epoch1) | *Arc+*<br>(Epoch2) | Paired mean difference<br>Epoch2 minus Epoch1 (95% CI) |
| --- | --- | --- | --- |
| CC | 0.012±0.01 | 0.035±0.01 | 0.023 [0.011, 0.030] |
| A/A | 0.361±0.04 | 0.346±0.04 | -0.015 [-0.031, 0.001] |
| A/B | 0.390±0.07 | 0.327±0.08 | -0.063 [-0.078, -0.053] |
| A25/B | 0.305±0.06 | 0.371±0.04 | 0.067 [-0.019, 0.139] |
| A/B25 | 0.347±0.07 | 0.109±0.05 | -0.238 [-0.298, -0.205] |
| A30 | 0.279±0.03 | 0.132±0.10 | -0.147 [-0.259, -0.067] |

| Group | Similarity<br>score | Paired mean difference<br>'Group' minus CC (95% CI) |
| --- | --- | --- |
| CC | -0.027±0.03 | 0 |
| A/A | 0.432±0.05 | 0.459 [0.412, 0.505] |
| A/B | 0.265±0.02 | 0.291 [0.256, 0.319] |
| A25/B | 0.255±0.02 | 0.278 [0.244, 0.309] |
| A/25B | 0.456±0.14 | 0.483 [0.357, 0.598] |
| A30 | 0.575±0.11 | 0.602 [0.502, 0.702] |

#### 3.1.1. Behavior

##### 3.1.1.1. Locomotion during the two epochs

We assessed the distance traversed during the first (Epoch1) and last (Epoch2) 5 minutes of the 30-minute experimental period matching the epochs used for IEG measurement. Two-way RM ANOVA on the distance traversed consisted of the ‘Group’ factor (A/A, A30, A/B25, A25/B, A/B) and the repeated measures factor ‘Epoch’ (1 and 2). The ANOVA revealed significant effects of Group (F_4,15_=19.50, p<0.0001), Epoch (F_1,15_=137.0, p<0.0001) and the Group×Epoch interaction (F_4,15_=36.67, p<0.0001) on the distance traversed. Šídák’s multiple comparisons test found no intergroup differences in Epoch1 as would be expected due to all experimental groups having the same conditions in the first epoch. In Epoch2, sharp differences were revealed between the groups (**Fig. 3**). Specifically, rats in the A30 and A/B25 groups traversed much shorter distances than the other groups (adjusted p<0.0001 for all comparisons) but did not differ from each other (p=0.2939).

**Figure 3.**
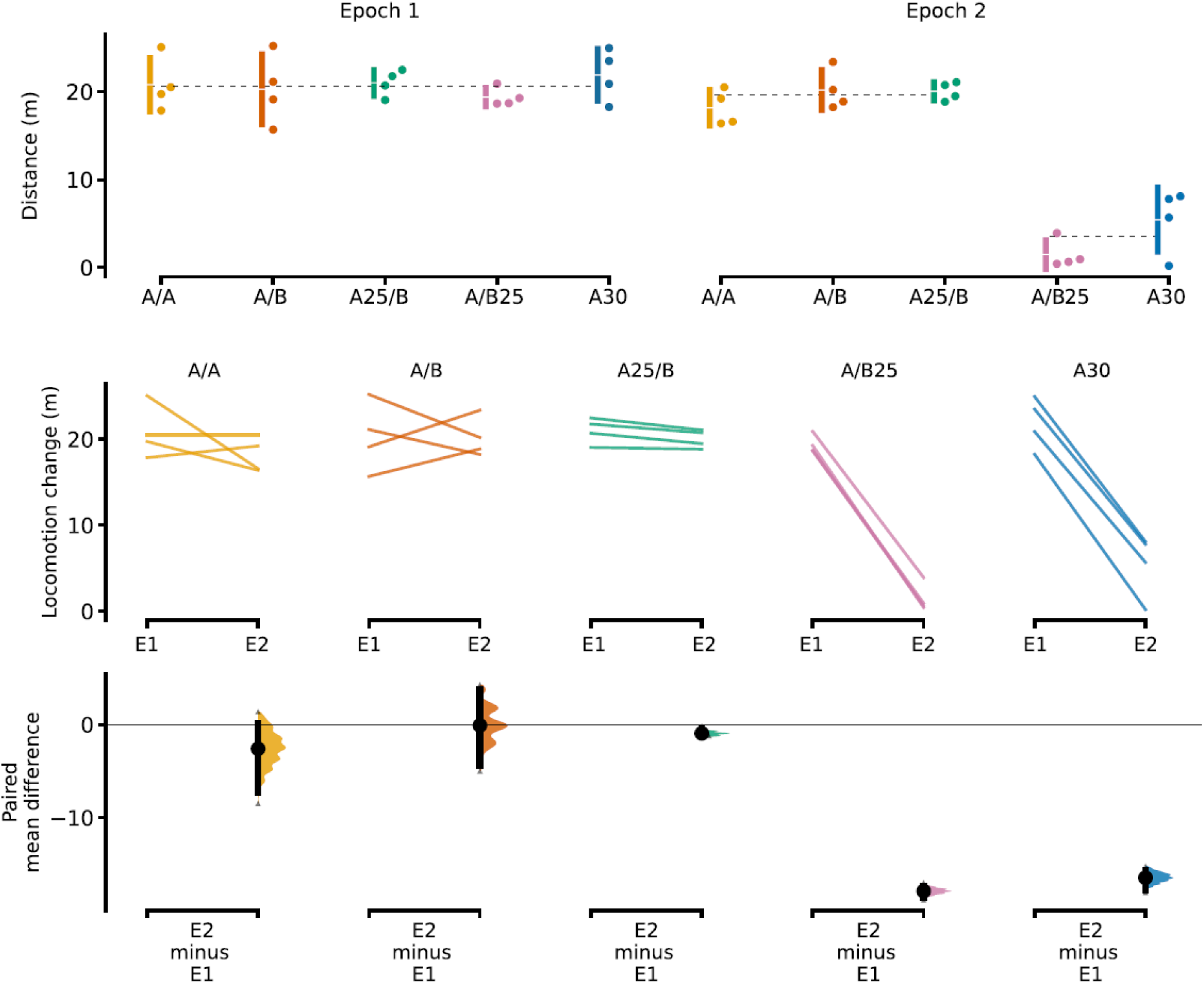
Locomotion during Epochs 1 and 2. **Top:** Total distance traversed by rats during Epoch1 (left) and Epoch2 (right), i.e. when IEG expression was induced. Each epoch lasted 5 minutes. Dashed horizontal lines show mean values: for Epoch1 the mean of all active groups (20.7±2.5); for Epoch2 the mean of recently transferred groups (19.3±1.9, left) and groups with continuous presence in final context (3.4±3.4, right). Vertical lines are standard deviations with the gap representing the mean value of each group. Epoch1: no significant differences between groups; Epoch2: ‘Recently transferred groups’ (A/A, A/B, A25/B) > ‘Continuous presence groups’ (A/B25, A30), p<0.0001. **Middle:** Raw data of locomotion change between the two epochs for each group, each paired set (Epoch1 to Epoch2) is connected by a line. **Bottom:** Dots represent the mean difference of locomotion between the two episodes; vertical error bars are 95% confidence intervals with bootstrapped sampling distribution. N=4 in each group.

Although contexts A and B had different shapes and slightly different areas, the differences in locomotion between epochs for groups A/A, A/B, and A25/B were negligible. In contrast, locomotion during the second epoch was greatly reduced in groups A/B25 and A30. Crucially, the analysis confirmed that the key factor was recency of transfer relative to Epoch2, and not physical properties of environments A and B. Animals transferred right before the Epoch2 (A/A, A25/B, and A/B) were much more active than animals already present in the environment (A30, A/B25) due to habituation of the initial exploration.

#### 3.1.2. Expression of *Homer1a* and *Arc*

##### 3.1.2.1. Ensemble sizes

Two-way RM ANOVA composed of ‘Group’ factor (CC, A/A, A/B, A25/B, A/B25, A30) and repeated measures factor ‘Epoch’ (the proportions of *Homer1a+* and *Arc+* nuclei representing ensemble activity during Epochs 1 and 2, respectively) revealed highly significant effects of Group (F_5,17_=24.40; p<0.0001), Epoch (F_1,17_=21.30; p=0.0002), and a Group×Epoch interaction (F_5,17_=12.07; p<0.0001) on the size of IEG ensembles.

Šídák’s multiple comparisons test for Epoch1 showed that all experimental groups had higher proportions of *Homer1a+* cells than the CC group (all p<0.0001), but no significant differences were found between the experimental groups in Epoch1. For Epoch2, groups A/A, A/B, and A25/B had more *Arc+* neurons than the CC group (all p<0.0001). Groups that were not recently transferred between environments (A/B25, A30) had proportions of *Arc+* cells significantly smaller compared to A/A (p<0.0001), A/B (p=0.0005), and A25/B (p<0.0001), and not significantly different from CC animals (A/B25 p=0.7969, A30 p=0.4129) nor from each other (p>0.9999). The ensemble size change between epochs was statistically significant (*Homer1a*>*Arc*) only in groups with continuous presence (A/B25, A30), p<0.05.

Proportions of IEG+ cells and ensemble size change are shown in **Fig. 4** Representative images of *Homer1a* and *Arc* RNA in situ hybridization are shown in **Fig. 5**.

**Figure 4.**
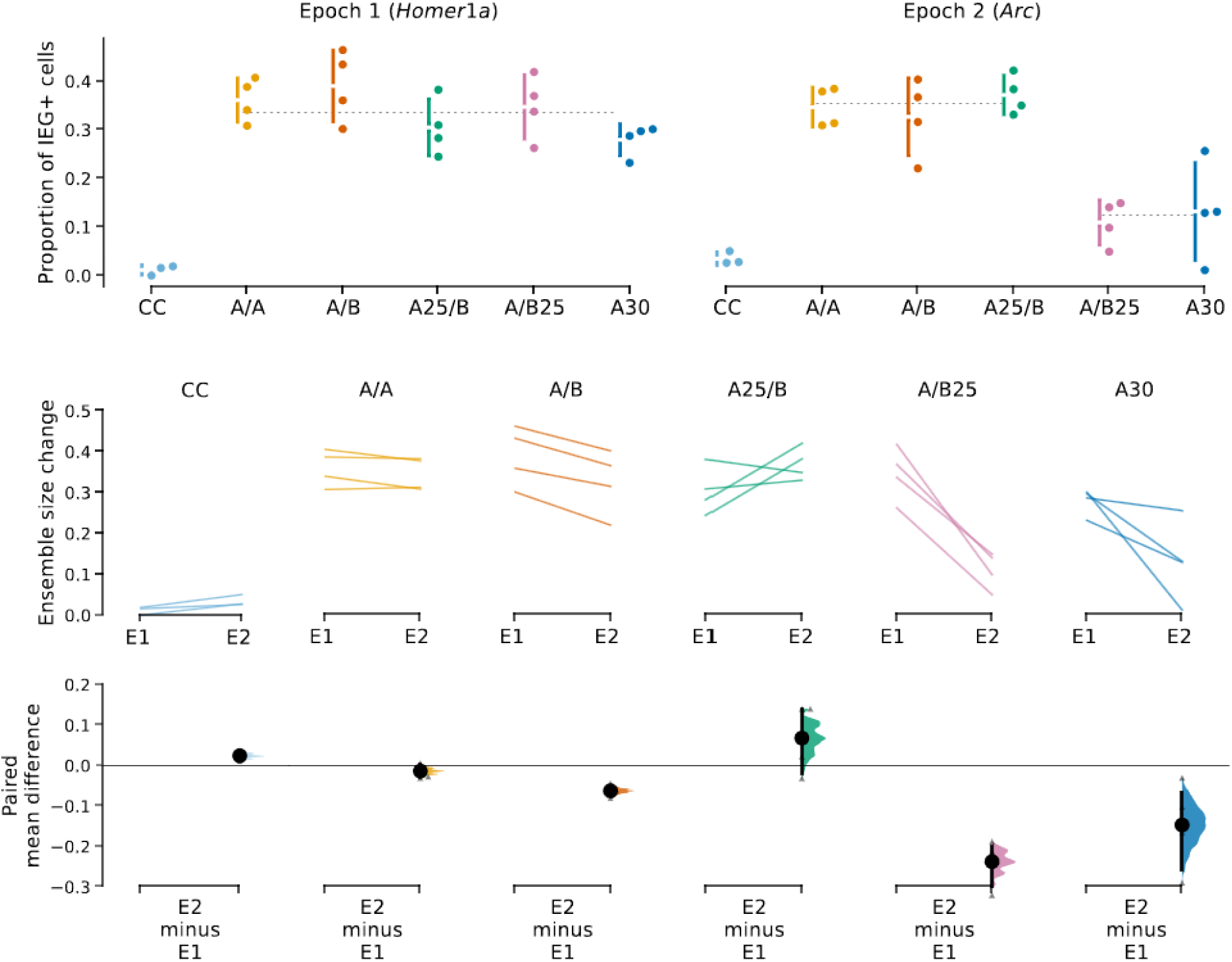
Homer1a and Arc ensemble size and dynamics. **Top:** Ensemble size marked as the proportion of cells expressing *Homer1a* (left, Epoch1) and *Arc* (right, Epoch2). Dashed horizontal lines display mean values: for Epoch1 of all active animals (*Homer1a* 0.336±0.06); for Epoch2 the mean of recently transferred groups (0.348±0.05, left) and groups with continuous presence in final context (0.121±0.07, right). Vertical lines are standard deviations with the gap representing the mean value of each group. Epoch1: (all active groups) > CC (p<0.0001); no significant differences between active groups. Epoch2: Recently transferred (A/A, p<0.0001; A/B, p<0.0005; A25/B, p<0.0001) > Continuous presence (A/B25, A30) = CC. **Middle:** Ensemble sizes at two epochs displaying individual rats in each group, each paired set is connected by a line (Epoch1 to Epoch2). **Bottom:** Estimation of ensemble size change. Dots represent mean differences in ensemble sizes between the two epochs; vertical error bars are 95% CIs with bootstrapped sampling distribution. N=4 in each group, except for CC where N=3.

**Figure 5.**
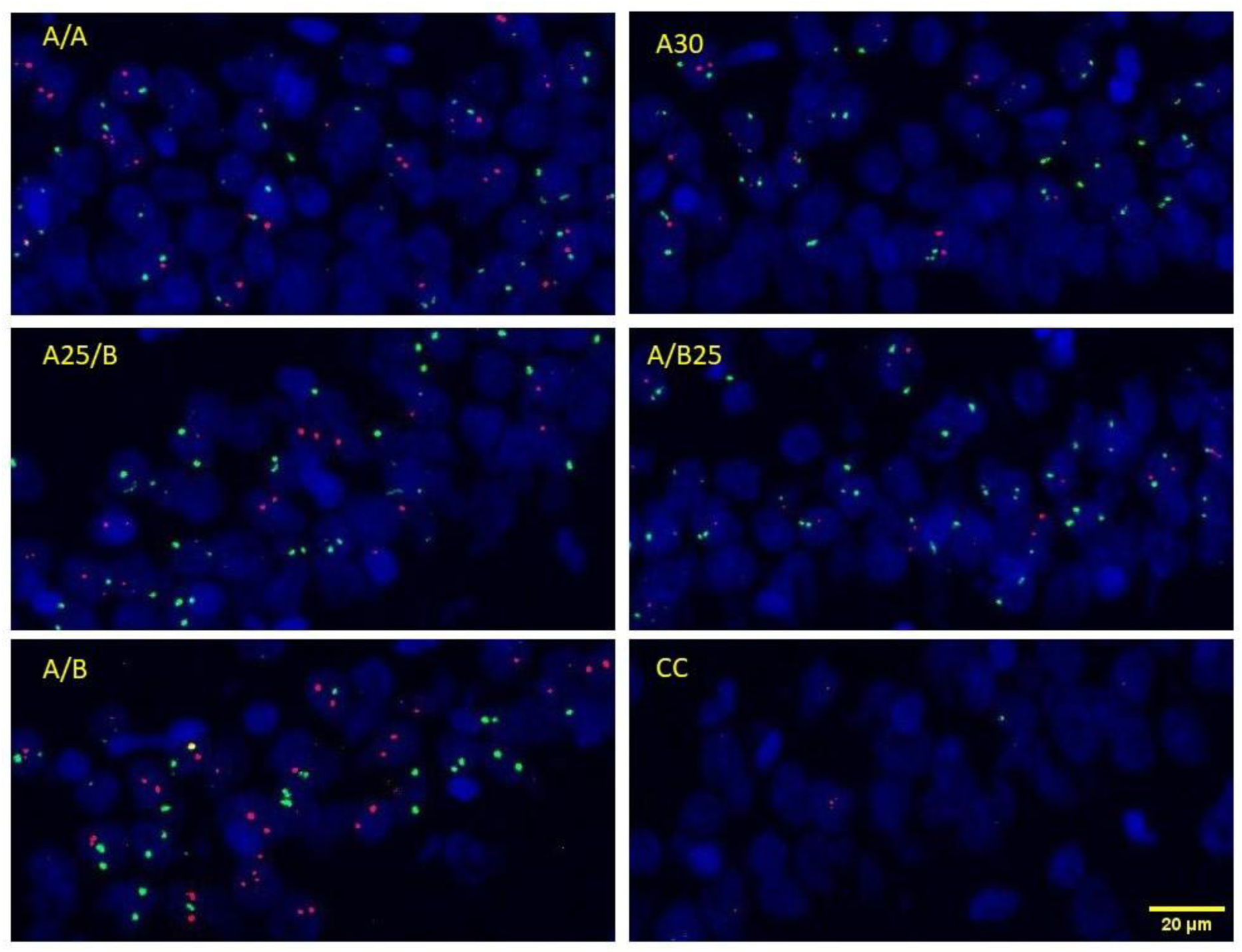
Representative images of Arc/Homer1a catFISH. *Homer1a* (green) and *Arc* (magenta) mark neurons active during Epochs 1 and 2, respectively. Neuronal (larger, semi-circular) and glial (smaller, denser and less circular) nuclei are counterstained with DAPI (blue). Note the high proportions of *Arc+* nuclei in A/A, A25/B, and A/B (left panels) contrast with the low proportions of *Arc+* nuclei in A30 and A/B25 (top 2 right panels) and only a few positive neurons present in cage controls (CC, right bottom panel). The red and green signals in these images were enhanced for the sake of illustration.

##### 3.1.2.2. Ensemble similarity

One-way ANOVA on similarity scores found significant differences among the groups (F_5,17_=23.89, p<0.0001). We used Holm-Šídák’s multiple comparison test to compare differences between the groups. All experimental groups displayed significantly higher similarity scores compared to CC (**Fig. 6**): A/A (p<0.0001), A/B (p=0.0018), A25/B (p=0.0026), A/B25 and A30 (p<0.0001). The test also confirmed higher similarity in A/A compared to A/B (p=0.0436) and A25/B (p=0.0313). The A/B and A25/B groups (recent transfer to B) did not differ from each other (p=0.8962). Groups A/B25 and A30, which remained in the same environment for Epoch2, were also not different from each other (A/B25 vs A30 p=0.1453), but they had significantly higher similarity scores compared to recently transferred groups: specifically, A/B25 compared to A/B (p=0.0248) and A25/B (p=0.0168); and A30 compared to A/B (p=0.0004) and A25/B (p=0.0003).

**Figure 6.**
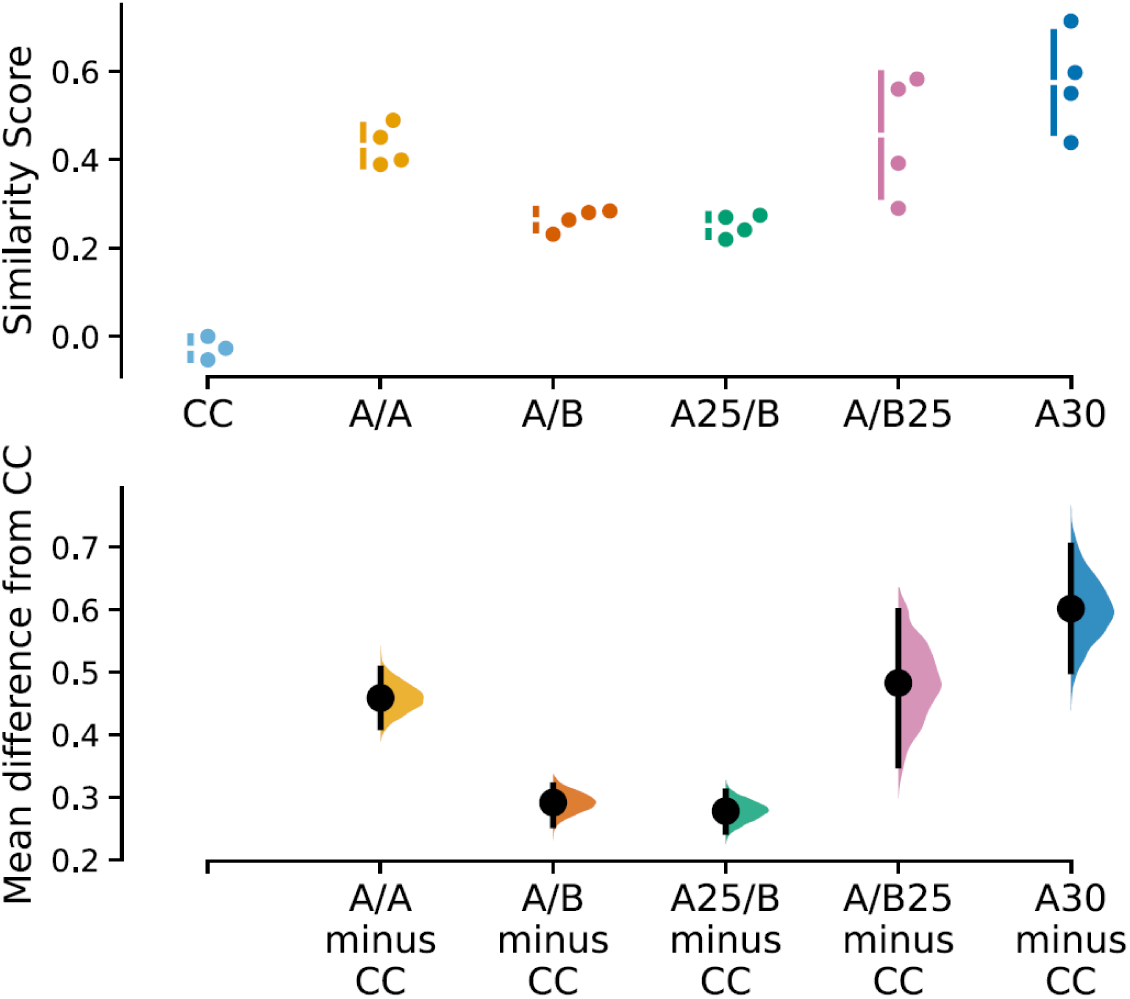
Similarity of ensembles active in the two epochs. **Top:** Dots represent individual animals. Vertical error bars are standard deviations with the gap representing the mean value of each group. **Bottom:** Each experimental group is compared to CC. Dots represent mean differences from the shared control of the CC group; vertical error bars are 95% CIs with bootstrapped sampling distribution. For p-values see section 3.1.2.2. N=4 in each group, except for CC where N=3.

### 3.2. Experiment 2 - Effect of robot avoidance and extinction on the size and overlap of IEG-expressing neuronal ensembles

Overview of effect sizes for Experiment 2 is provided in **Tab. 3**.

**Table 3.** Effect Sizes for Experiment 2. Locomotion, Shocks during Avoidance Training, Ensemble sizes, and Ensemble overlap.

| Group | Epoch1 | Epoch2 | Paired mean difference<br>Epoch2 minus Epoch1 (95% CI) |
| --- | --- | --- | --- |
| CC | x | x | x |
| nsSS | 15.4 ± 5.5 | 15.0 ± 10.4 | -0.48 [-7.29, 5.59] |
| SS | 13.2 ± 6.9 | 11.6 ± 8.2 | -1.64 [-4.51, 3.52] |
| SM | 13.1 ± 8.8 | 12.2 ± 9.2 | -0.985 [-5.86, 5.48] |
| MS | 17.0 ± 6.9 | 19.6 ± 8.8 | 2.58 [-1.13, 5.13] |
| MM | 13.4 ± 5.0 | 12.1 ± 6.6 | -1.38 [-5.86, 3.05] |

| Group | Day1 minus Day2 | Day2 minus Day3 | Day3 minus Day4 |
| --- | --- | --- | --- |
| CC | x | x | x |
| nsSS | 56.5 [-64.2, 156] | 82.5 [-30.5, 266] | -45.8 [-197, 26.4] |
| SS | 3.4 [-33.6, 92.8] | -29.6 [-81.9, -4.0] | -18.9 [-34.5, -12.2] |
| SM | -12.2 [-63.0, 15.2] | -12.0 [-36.8, 4.9] | -11.2 [-18.1, -4.62] |
| MS | -27.0 [-63.4, 8.3] | -23.9 [-52.4, 2.8] | -13.2 [-45.6, 9.4] |
| MM | -14.5 [-37.6, 19.8] | -40.2 [-84.2, -16.1] | -29.2 [-40.2, -15.2] |

***Ensemble size: Proportion IEG+ cells (mean, std)***
| Group | Homer1a+<br>(Epoch1) | Arc+<br>(Epoch2) | Paired mean difference<br>Epoch2-Epoch1 (95% CI) |
| --- | --- | --- | --- |
| CC | 0.054 ± 0.04 | 0.057 ± 0.03 | -0.003 [-0.056, 0.033] |
| nsSS | 0.393 ± 0.05 | 0.215 ± 0.07 | -0.178 [-0.218, -0.129] |
| SS | 0.317 ± 0.06 | 0.182 ± 0.06 | -0.135 [-0.170, -0.092] |
| SM | 0.344 ± 0.09 | 0.228 ± 0.08 | -0.116 [-0.164, -0.054] |
| MS | 0.349 ± 0.13 | 0.227 ± 0.07 | -0.131 [-0.184, -0.080] |
| MM | 0.366 ± 0.09 | 0.227 ± 0.08 | -0.140 [-0.210, -0.065] |

Overall mean of all active animals in Epoch1 = 0.336 ± 0.06 ***Ensemble overlap (mean, std)***
| Group | Similarity score | Paired mean difference<br>'Group' minus CC (95% CI) |
| --- | --- | --- |
| CC | 0.055 ± 0.10 | 0 |
| nsSS | 0.435 ± 0.07 | 0.380 [ 0.296, 0.453] |
| SS | 0.401 ± 0.21 | 0.346 [0.186, 0.491] |
| SM | 0.396 ± 0.09 | 0.340 [0.252, 0.419] |
| MS | 0.454 ± 0.11 | 0.399 [0.305, 0.493] |
| MM | 0.389 ± 0.07 | 0.334 [0.255, 0.409] |

Overall mean of all active animals = 0.4215 ± 0.116

#### 3.2.1. Behavior

##### 3.2.1.1. Robot Avoidance Learning

All rats in the experimental groups (SS, SM, MS, MM) learned to avoid the robot as shown by the gradual decrease in the number of shocks delivered on each training day (**Fig. 7**). In contrast, the active controls (nsSS) showed no decrease from day 1 to 4. Two-way RM ANOVA performed on the training days found significant effects of Days (F_3,105_=12.66; p<0.0001) and Groups (F_4,35_=7.392; p<0.0002), but not the interaction (F_12,105_=1.745; p=0.0571). Holm-Šidák’s test on the main effect of Days revealed that rats significantly reduced the number of received shocks during the first three days of training: day 1 vs 2 p<0.0001; day 2 vs 3 p=0.0017. No statistically significant difference was observed between days 3 and 4 (p=0.3534).

**Figure 7.**
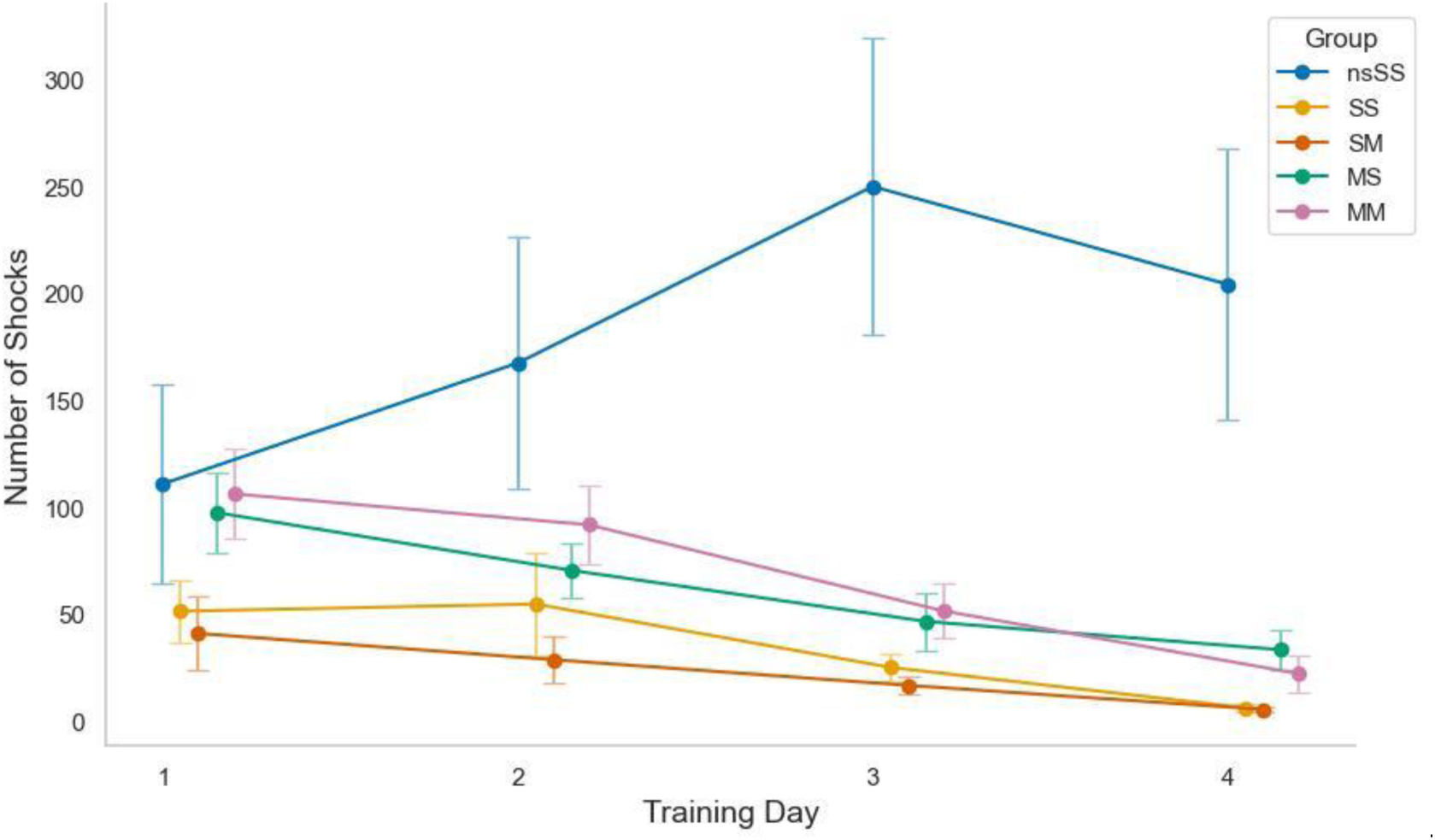
Robot Avoidance Training in Experiment 2. Each training session lasted 20 min/day for four days. Rats in the SS and SM groups were trained to avoid a stationary robot, the MS and MM groups were trained with a moving robot. All trained groups gradually reduced the number of shocks received over the training period, whereas the no-shock active controls nsSS did not (virtual shocks shown). The overall mean of trained groups on day 1 was 74.0±10.0 SEM, and 16.8±3.7 SEM on day 4. Day1>Day2 (p<0.0001); Day2>Day3(p=0.0017); Day3 vs Day4 (n.s.). Error bars are SEM. N=8 for each group.

##### 3.2.1.2. Locomotion during the two epochs of the test session

The total path traversed during the two epochs (0-5 min, 25-30 min) of the test session (**Fig. 8**) did not significantly differ between the active groups F(_4, 35_)=1.095; p=0.3742 or the two epochs F(_1, 35_)=0.1011; p=0.7524; no significant interaction was present F(_4, 35_)=0.4056; p=0.8033.

**Figure 8.**
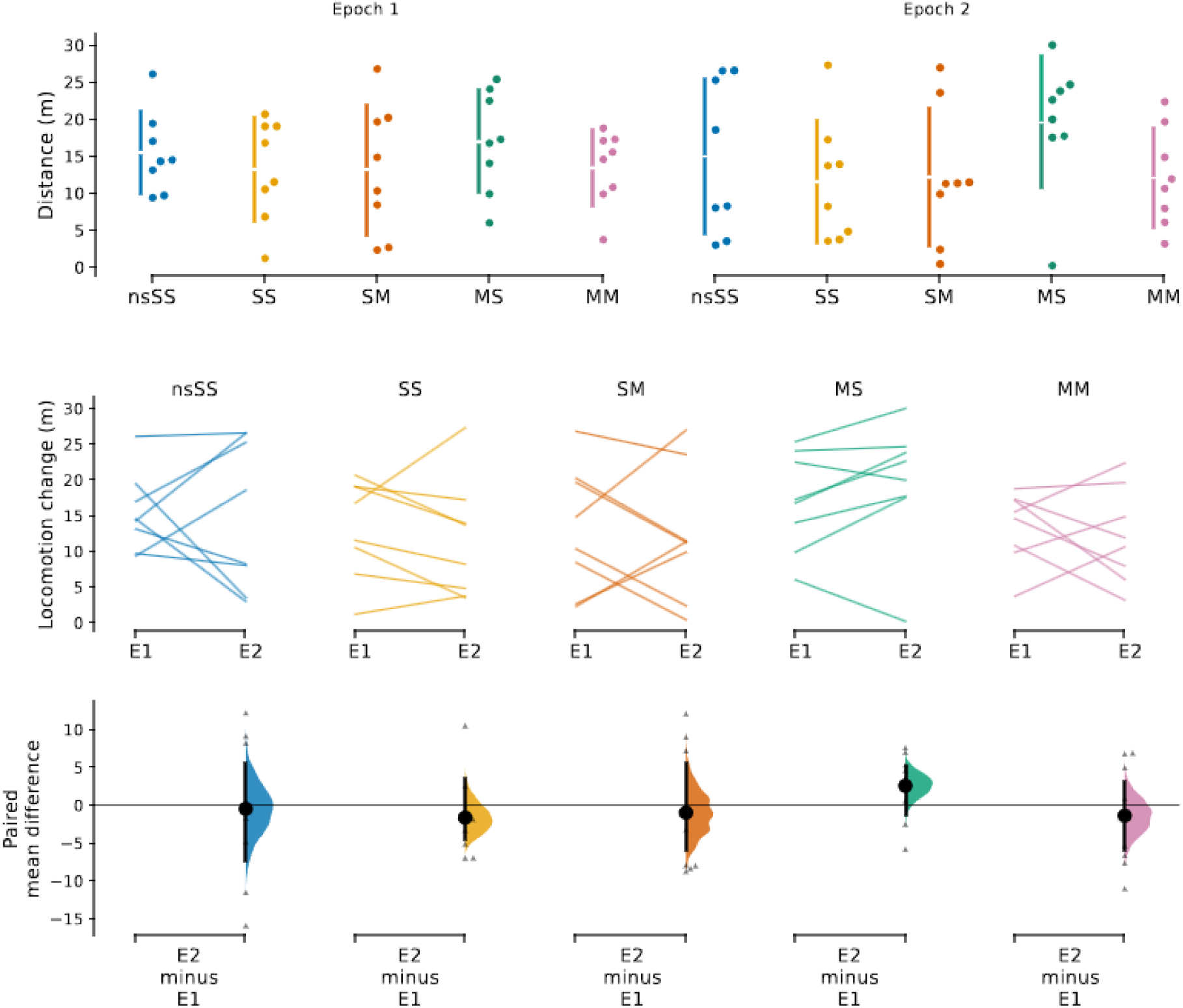
Locomotion during Epochs 1 and 2. **Top:** Distance in meters traversed during the first (left, Epoch1) and the last (right, Epoch2) 5 minutes of the continuous 30-minute testing session. Dots represent individual rats; error bars are standard deviations, the gap in the error bar marks the mean value of each group. No significant differences between groups in either epoch. **Middle:** Comparison of locomotion between the two epochs, raw data of individual rats, each paired set of Epoch1 and Epoch2 is connected by a line. **Bottom:** Paired mean differences (m) for each group. Dots represent mean difference of locomotion between the two epochs; vertical error bars are 95% CIs with bootstrapped sampling distribution. N=8 in each group.

#### 3.2.2. Expression of *Homer1a* and *Arc*

##### 3.2.2.1. Ensemble size

*Homer1a+* cells marked ensembles active in the first five minutes (Epoch1) of the 30-minute continuous test session, while Arc+ cells marked ensembles active in the last five minutes (Epoch2) of the test session. The two-way RM ANOVA on the proportions of IEG-expressing neurons with ‘Group’ factor (CC, nsSS, SS, MS, SM, MM) and repeated measures on the Epoch factor (*Homer1a*, *Arc*) revealed significant effects of Group (F_5,42_=17.75; p<0.0001), Epoch (F_5,42_=98.80; p<0.0001), and Group×Epoch interaction (F_5,42_=4.63.75; p<0.0019). Šidák’s post-hoc test confirmed that passive caged controls had markedly smaller IEG+ ensembles than all other groups: in Epoch1 CC vs. all other groups p<0.0001, in Epoch2 CC vs. nsSS, MS, SM, MM p<0.001 and CC vs. SS p=0.0138 (Fig. 9). However, no statistically significant differences between active groups (nsSS, SS, MS, SM, MM) were present in either epoch.

**Figure 9.**
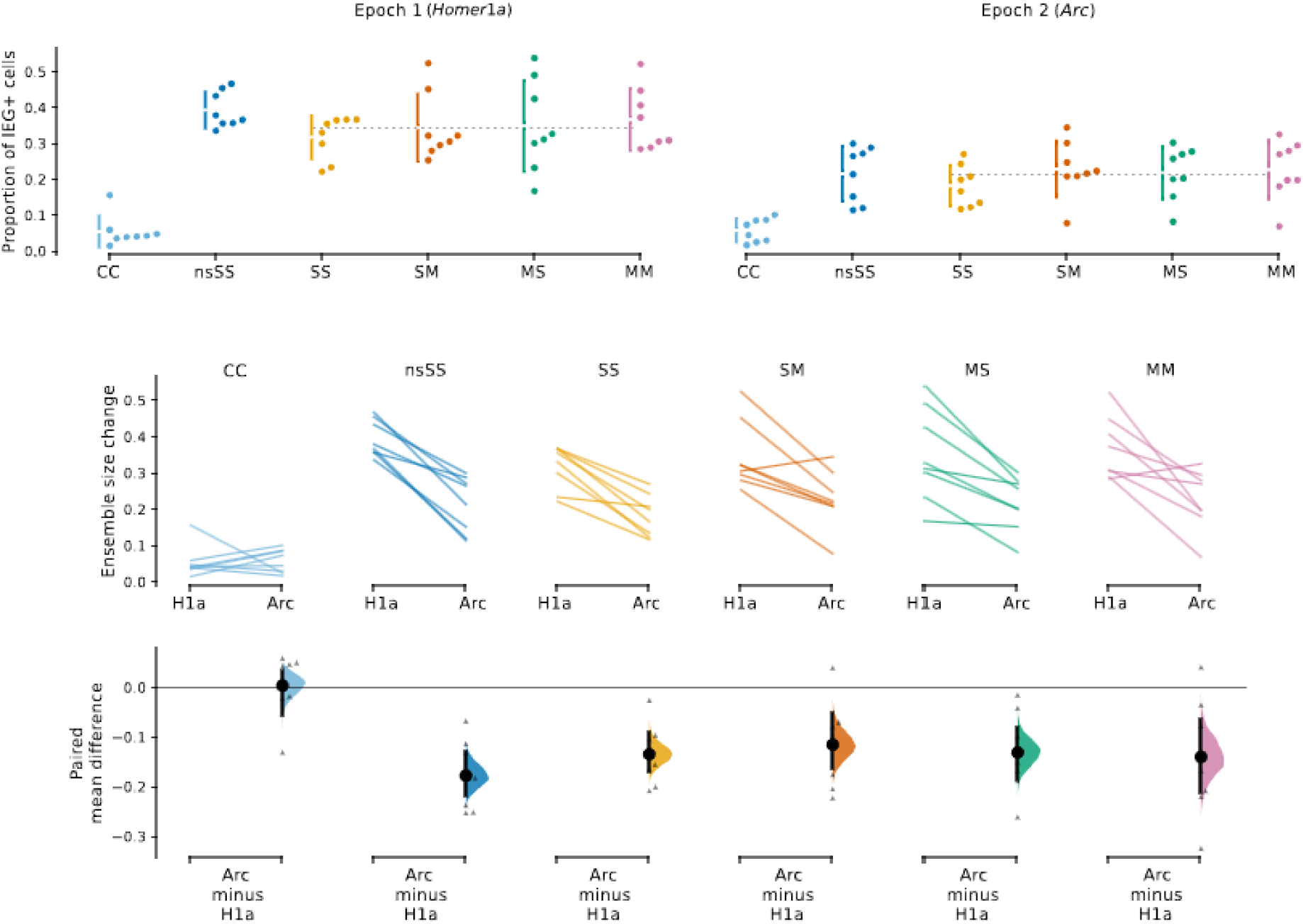
Proportion of Homer1a and Arc-positive cells and ensemble size change between epochs. **Top:** Proportion of cells expressing *Homer1a* (left, Epoch1) and *Arc* (right, Epoch2) during the 30-minute test session, no shocks were delivered during the test. Dots represent individual animals; vertical error bars are standard deviations of each group with the gap showing the mean value. Dashed horizontal line shows the mean value of IEG expression of all trained animals (nsSS excluded, *Homer1a* 0.354±0.09 std, *Arc 0.*214±0.07 std). Active groups > CC (Epoch1 p<0.0001; Epoch2 p<0.05); Epoch1 > Epoch2 for Active groups (p<0.0001); no significant differences between active groups were found in either epoch. **Middle:** Paired mean differences for comparison of IEG ensemble size in each epoch showing ensemble sizes and change for individual rats in each group. Each paired set (Epoch1 to Epoch2) is connected by a line. **Bottom:** Paired mean difference of ensemble size between the epochs for each group. Dots represent mean differences; vertical error bars are 95% confidence intervals with bootstrapped sampling distribution. N=8 in each group.

##### 3.2.2.2. Ensemble overlap

To assess the overlap between ensembles active at the start (*Homer1a*, Epoch1) and the end (*Arc*, Epoch2) of the test session, we calculated similarity scores for each group. One-way ANOVA showed a statistically significant effect of Group: F(_5, 42_)=12.914; p=<0.0001) (Fig. 10). In line with previous reports (Kubik et al., 2014; Vazdarjanova and Guzowski, 2004), the similarity in the CC group was markedly lower than in the active groups. No statistically significant differences were present among the active groups. The congruence of training/test conditions ([SS, MM] vs. [MS, SM]) did not affect ensemble similarity.

**Figure 10.**
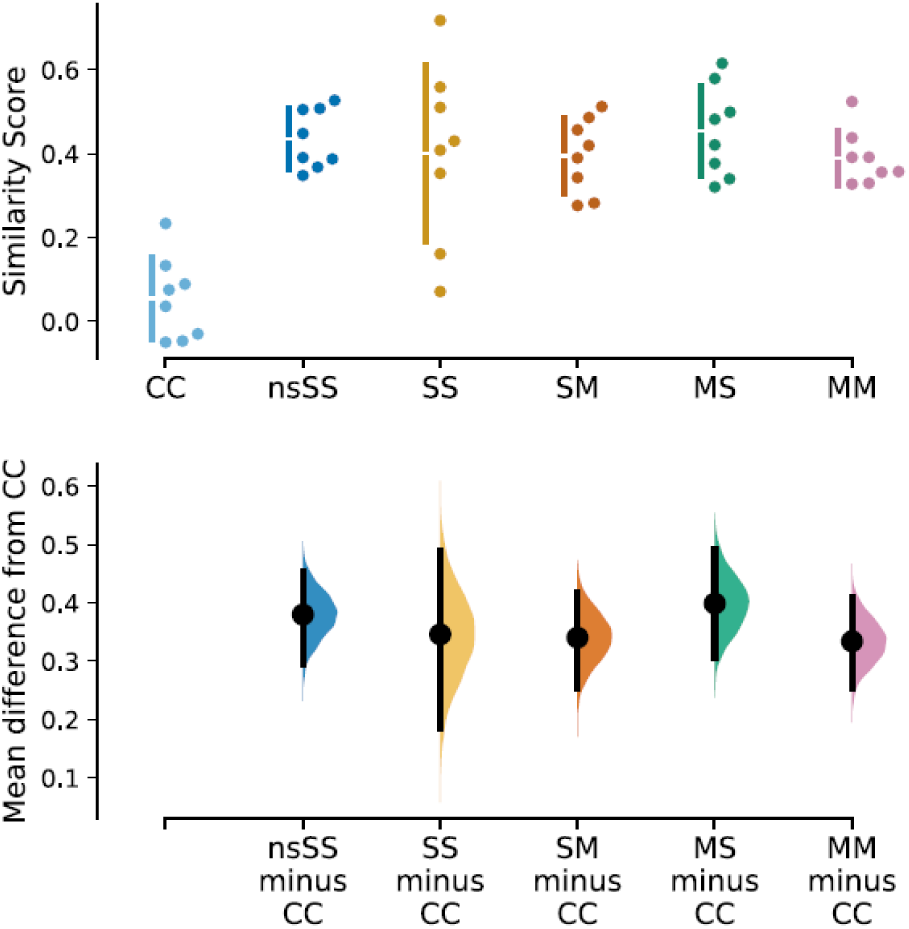
Similarity of ensembles active in the two epochs. The overlap of the IEG-expressing ensembles is indicated by a similarity score that is normalized for the cell population size. **Top:** Each dot represents a similarity score of a single animal. Vertical error bars are standard deviations with the gap representing the mean value of each group. ‘Active groups’ > CC (p=<0.0001); No significant differences between active groups. **Bottom:** Dots represent mean differences from the shared control of the CC group; vertical error bars are 95% confidence intervals with bootstrapped sampling distribution. N=8 in each group.

#### 3.2.3. Extinction of the Avoidance Response and its effect on *Homer1a/Arc+* ensembles

The paired mean differences between day 5 virtual and day 4 actual shocks accumulated at t=0-5 minutes (95% CI): SS 13.8 [1.38, 32.5], SM 31.5 [13.9, 54.9], MS -1.38 [-15.2, 14.2], MM 9.38 [-1.88, 41.0] (Fig. 11). All groups except MS received more virtual shocks during the first 5 minutes of the test than actual shocks on the previous day. However, some rats retained the avoidance response as their virtual shocks were lower, similar, or only slightly higher than on day 4, others clearly stopped avoiding the robot.

**Figure 11.**
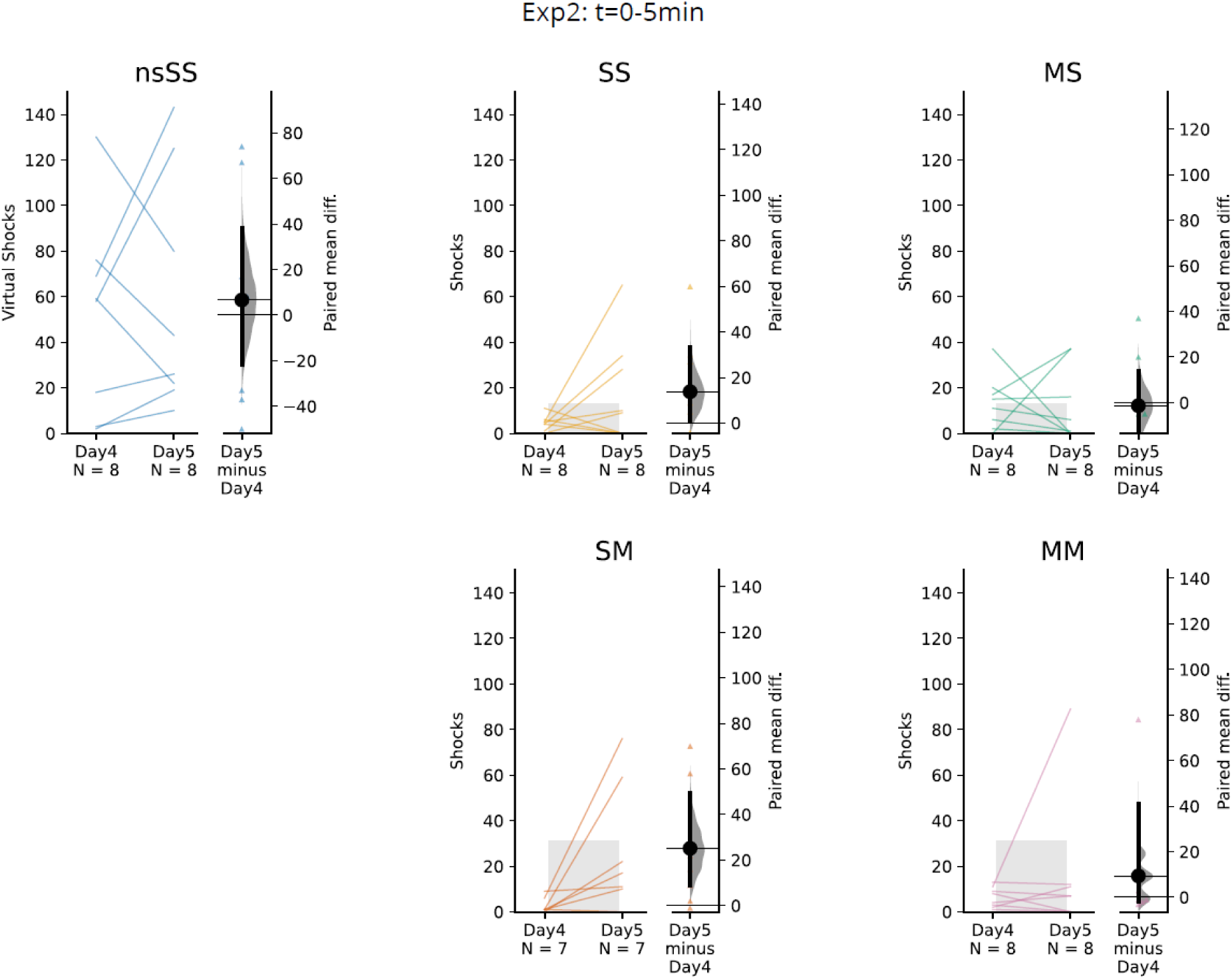
Comparison of actual and virtual shocks delivered by 5 minutes of the training day 4 and test day 5. Raw paired data of individual rats depicting the number of actual and virtual shocks received during the first 5 minutes of training on day 4 and test on day 5, respectively. The right portion of each plot shows 95% CI for paired mean differences of day 4 and day 5 for the whole group with distribution estimate. Gray bars represent one-sided tolerance intervals (TI, 99% proportion of population, 95% confidence level) later used as thresholds. TIs based on training day 4 data pooled from Experiments 2 and 3 at t=0-5min as described in Methods. S-5 TI applied to groups with a stationary robot during test (SS, MS); M-5 TI applied to groups with a moving robot during test (SM, MM).

The paired mean differences between virtual and actual shocks accumulated over 20 minutes (95% CI, day 5 minus day 4): SS 104 [47.2, 194], SM 132 [72.8, 198], MS 101 [56.2, 179], MM 39.2 [23.8, 66.9] (Fig. 12). The paired mean difference in untrained nsSS control group was close to zero at 5 minutes (6.62 [95% CI -21.9, 38.4]) and the smallest at 20 minutes (19.8 [95% CI -111, 153]).

**Figure 12.**
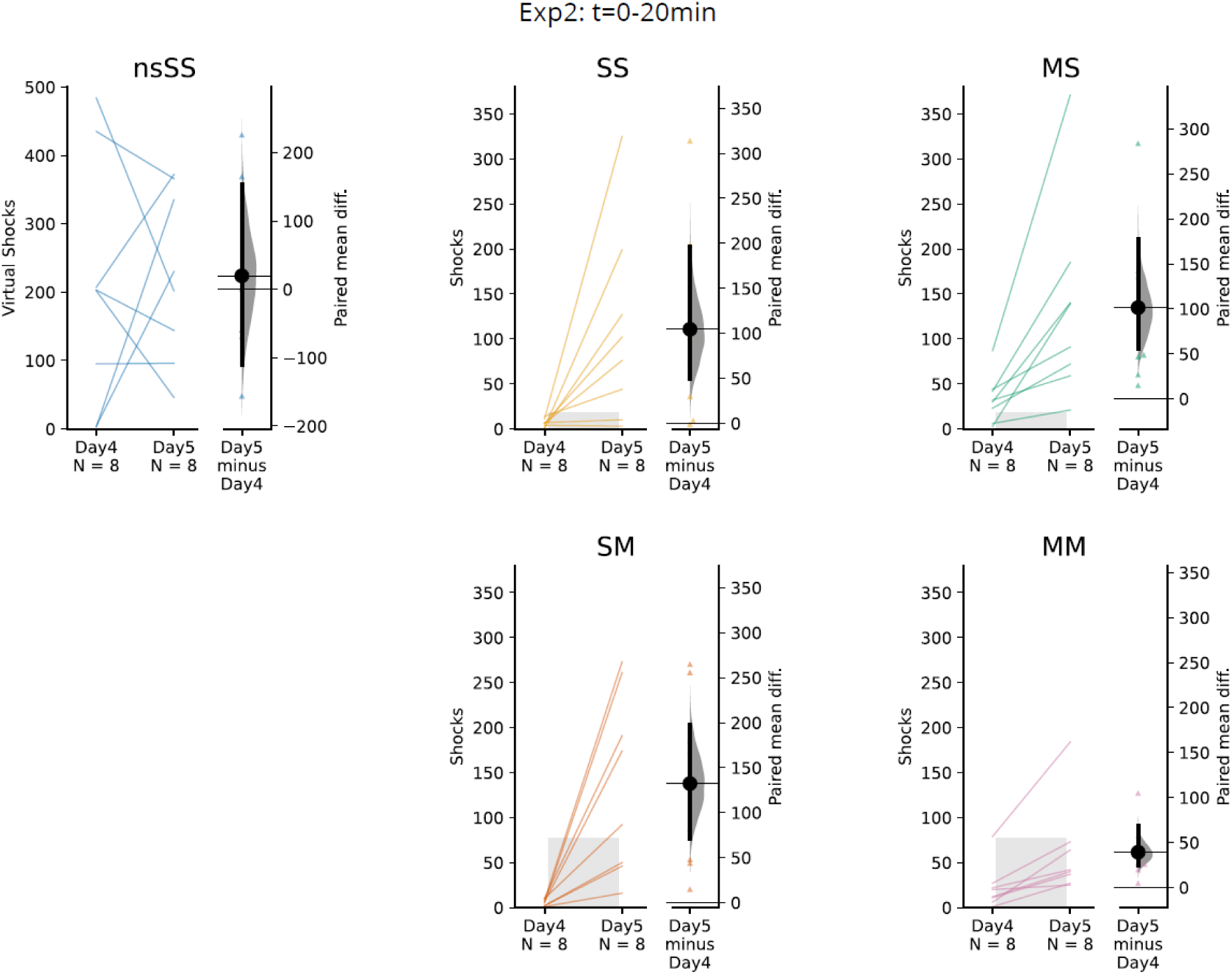
Comparison of actual and virtual shocks delivered at 20 minutes on training day 4 and test day 5. Raw paired data of individual rats depicting the number of actual and virtual shocks received during the first 20 minutes of training day 4 and test day 5, respectively. The nsSS active control group received no actual shocks during training; Note the different y-axis scale on the left-hand side in the nsSS group. The right portion of each plot shows 95% CI for paired mean differences of day 4 and day 5 with bootstrapped distribution estimates. Gray bars represent one-sided tolerance intervals (TI, 99% proportion of population, 95% confidence level) based on training day 4 data pooled from Experiments 2 and 3 at t=0-5min as described in Methods section 2.2.5. The S-20 TI threshold is applied to groups with robot stationary during the test (SS, MS); threshold TI M-20 is applied to groups with robot moving during the test (SM, MM).

As described in Methods (section 2.2.5.), as part of the *ex post* exploratory analysis we calculated one-sided tolerance intervals (TI, 99% proportion of population, 95% confidence level) for the number of shocks received on day 4 by animals pooled from Experiments 2 and 3. The TI calculated from groups with stationary robot (SS+SM) on day 4 at t=5min was 13 (threshold ‘S-5’), and at t=20min was 18 (threshold ‘S-20’); TI for groups with moving robot (MS+MM) on training day 4 at t=5min was 33 (threshold ‘M-5’), and at t=20min was 77 (threshold ‘M-20’). Applying the TIs as extinction thresholds (see section 2.2.5.) to trained animals in Exp2 (Fig. 13) showed that the numbers of rats with respect to avoidance/extinction in each epoch were: Av-Av N=4; Av-Ext N=10; Ext-Ext N=10; Unclear N=8.

**Figure 13.**
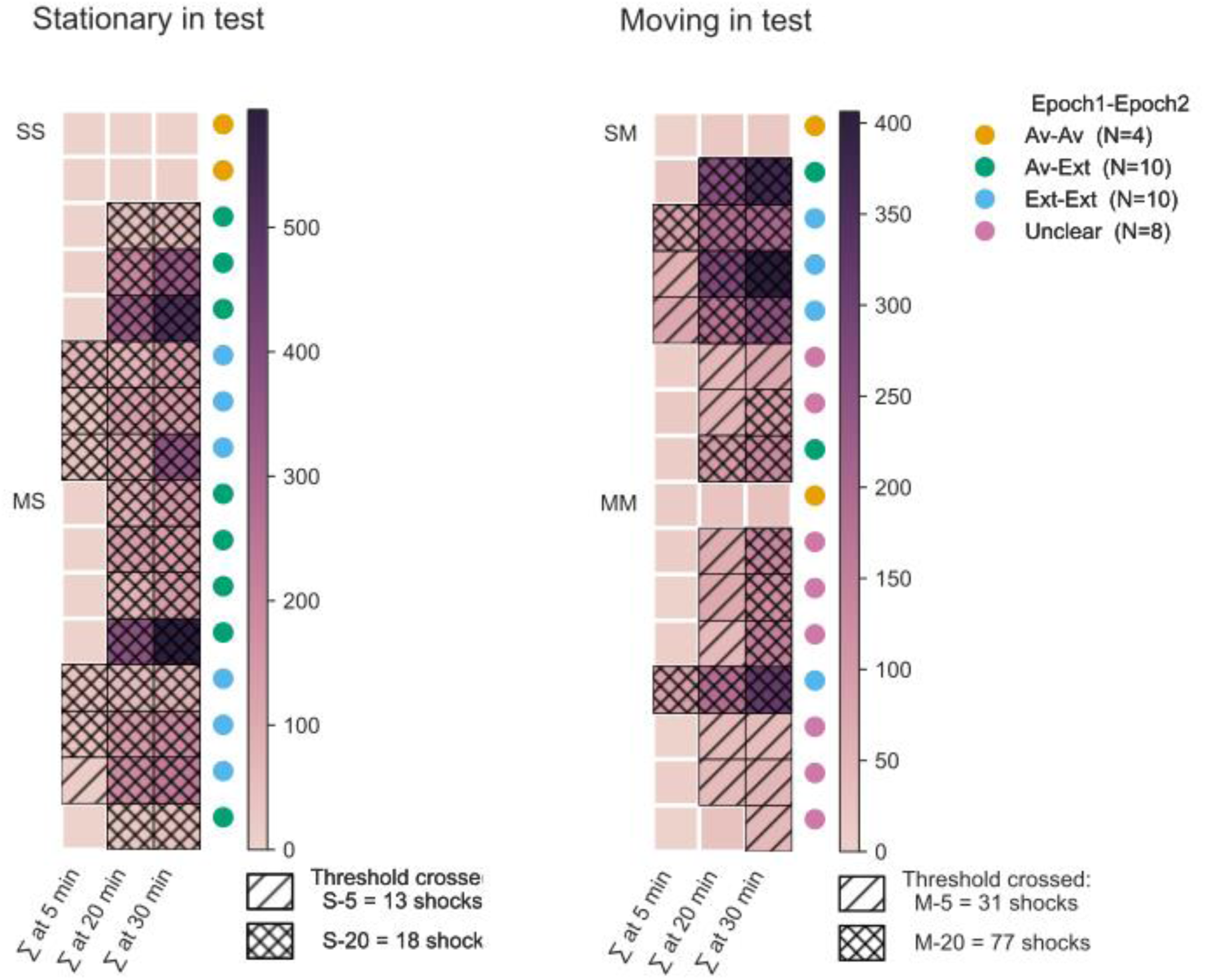
Avoidance and extinction groups in Experiment 2. Thresholds used to classify animals as ‘Av’ (avoidance) or ‘Ext’ (extinction) were based on one-sided tolerance intervals (99% proportion of population, 95% confidence level, TI) on the number of shocks received on training day 4 (Experiments 2 and 3 pooled); details on threshold calculation and application are specified in Methods section 2.2.5. Heatmap (colored squares) displays cumulative virtual shocks for each animal (y-axis) at a given time of the test (x-axis). The color bar for each heatmap is on the right-hand side of each heatmap. Hatchings mark different thresholds exceeded at a given time during the test. Color circles represent animals based on the presence of avoidance or extinction in Epochs 1 and 2.

The exploratory analysis showed that animals avoiding the robot in Epoch 1 (Av-Av, Av-Ext) had slightly smaller *Homer1a+* ensembles (0.313±0.08) than those in the nsSS (0.393±0.05) and Ext-Ext (0.382±0.11) groups, with a relative difference of ∼18%. Furthermore, the Ext-Ext group also displayed slightly larger *Arc+* ensembles (0.264±0.05) and a smaller ensemble size decline between epochs (-31%) compared to groups with late or no extinction (nsSS: 0.215±0.08, -45%; Av-Av: (0.225±0.10, -46%; Av-Ext: (0.177±0.06, -42%). This is in line with the notion that only limited learning occurred in the overtrained avoiding animals, whereas the Ext-Ext group rapidly learned the new contingencies (no shock). This extinction learning requires plasticity and IEG expression to accommodate new information and could delay the shrinking of IEG+ ensembles accompanying prolonged exploration **(Exp1)**. The proportions of *Homer1a+ and Arc+* cells, the relative change of ensemble size between epochs, and ensemble overlap (similarity scores) are shown in Fig. 14 and **Tab. 4**.

**Figure 14.**
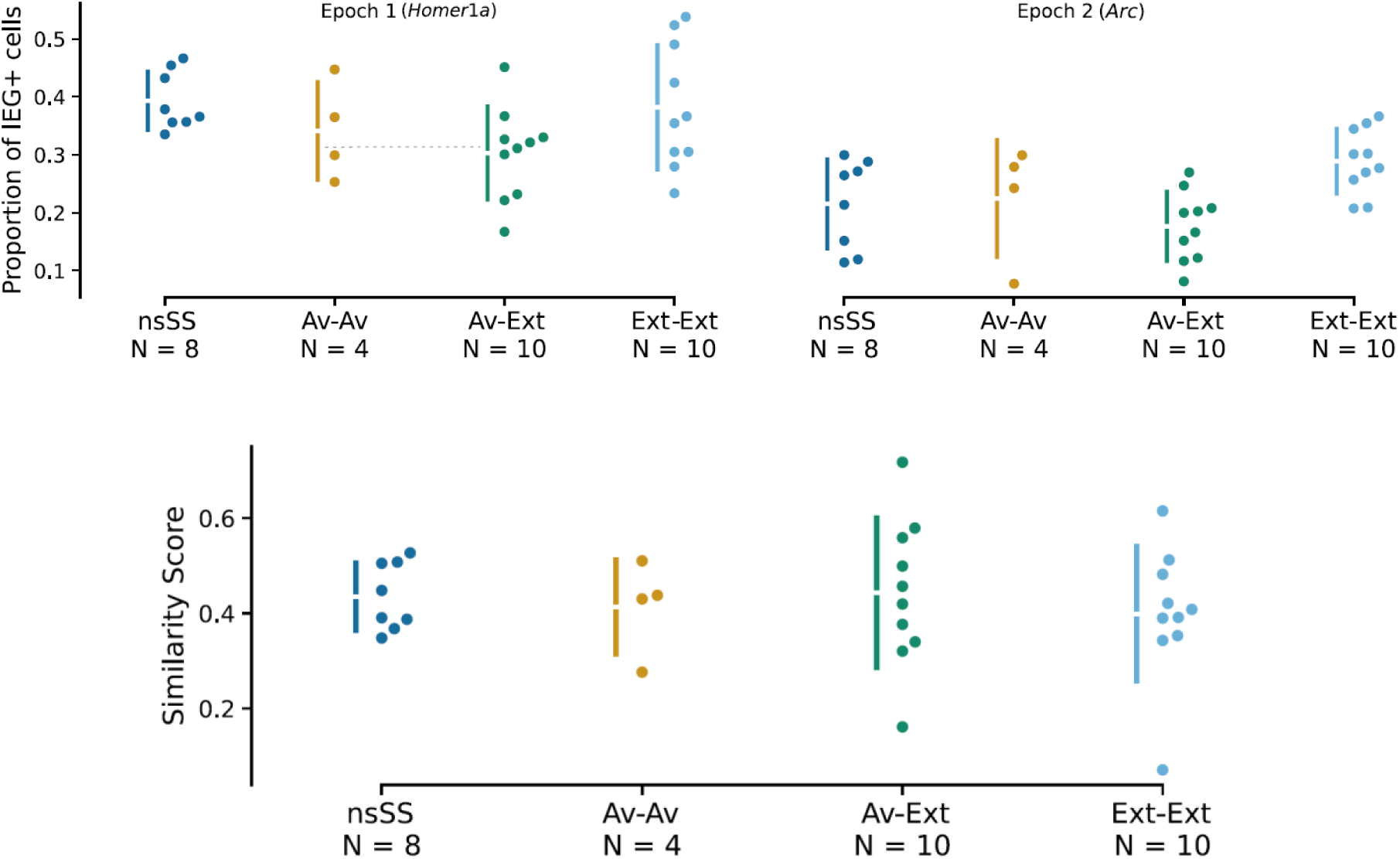
Homer1a/Arc Ensemble Size in each Epoch and Overlap relative to Avoidance and Extinction. **Top:** IEG expression for Epoch1 and 2 of the test session in relation to avoidance and extinction. Dots represent the proportions of *Homer1a* and *Arc*-positive cells in hippocampal CA1. Vertical error bars are standard deviations of each group with a gap showing the mean value. A dashed horizontal line shows the mean proportion of IEG+ cells in groups avoiding the robot in Epoch1 (*Homer1a*): 0.313±0.08 std; note that the average *Homer1a+* ensemble size in the Ext-Ext group is larger at 0.382±0.11. The Ext-Ext group also displayed the smallest ensemble size decrease between epochs of -31% (nsSS -45%; Av-Av -46%; Av-Ext -42%). **Bottom:** Similarity score measuring the overlap between IEG-positive ensembles activated at the two epochs in relation to avoidance and extinction in each of the epochs.

**Table 4.** IEG+ ensembles in Avoidance/Extinction groups. Ensemble size at each Epoch, Relative change of ensemble size, and Ensemble overlap.

**Ensemble Size: Proportion of IEG+ cells (mean, std)**
| Group | *Homer1a*+<br>(Epoch1) | *Arc*+<br>(Epoch2) | Relative change<br>(Epoch1 vs Epoch2) | N |
| --- | --- | --- | --- | --- |
| nsSS | 0.393 ± 0.05 | 0.215 ± 0.08 | -45% | 8 |
| Av-Av | 0.341 ± 0.08 | 0.225 ± 0.10 | -46% | 4 |
| Av-Ext | 0.303 ± 0.08 | 0.177 ± 0.06 | -42% | 10 |
| Ext-Ext | 0.382 ± 0.11 | 0.264 ± 0.05 | -31% | 10 |

| Group | Similarity score | N |
| --- | --- | --- |
| nsSS | 0.435 ± 0.07 | 8 |
| Av-Av | 0.414 ± 0.10 | 4 |
| Av-Ext | 0.443 ± 0.16 | 10 |
| Ext-Ext | 0.399 ± 0.14 | 10 |

**Table 5:**
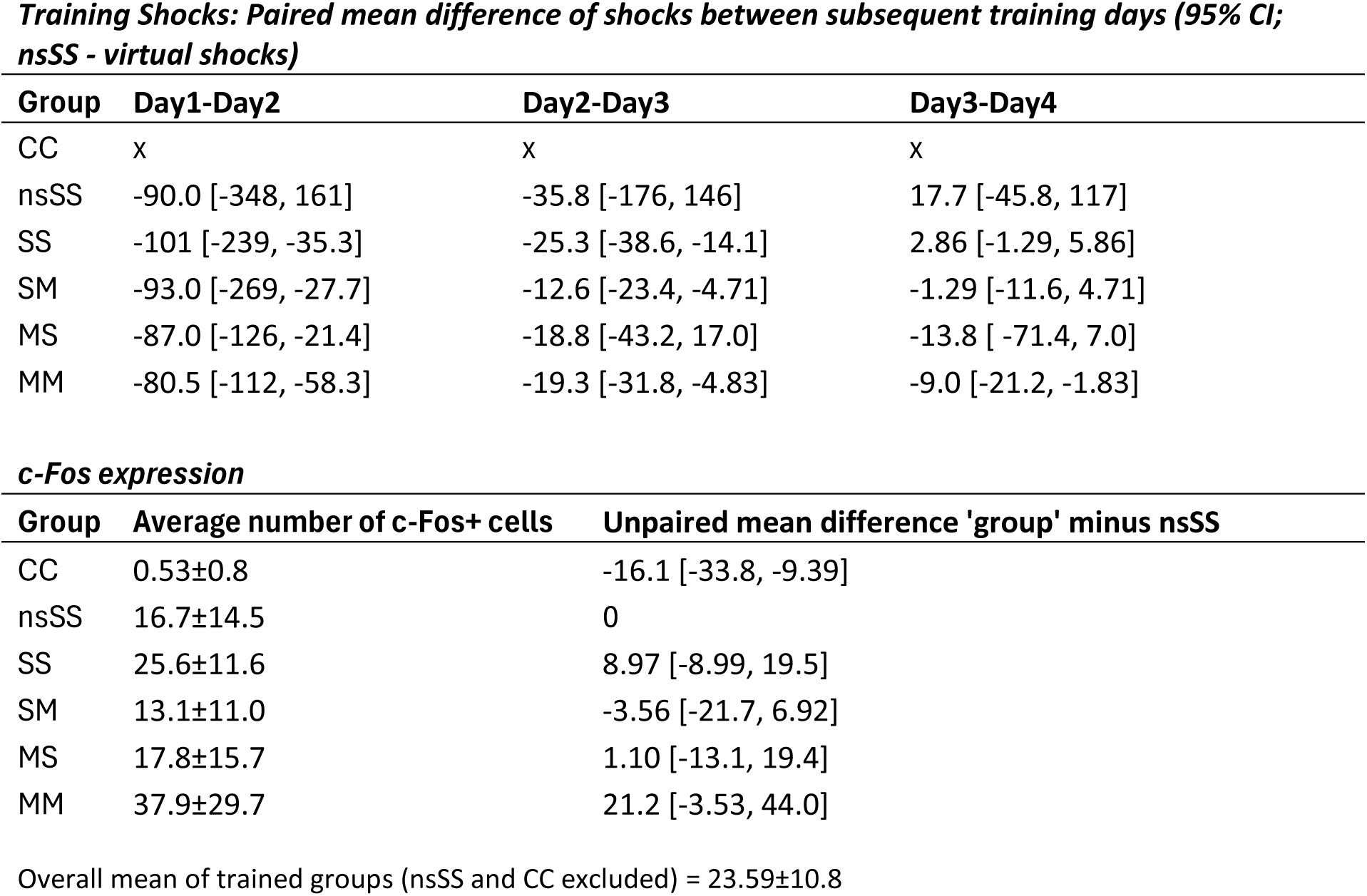
Effect Size Estimates for Experiment 3. Shocks during training, and c-Fos expression during test.

**Training Shocks: Paired mean difference of shocks between subsequent training days (95% CI; nsSS - virtual shocks)**
| Group | Day1-Day2 | Day2-Day3 | Day3-Day4 |
| --- | --- | --- | --- |
| CC | x | x | x |
| nsSS | -90.0 [-348, 161] | -35.8 [-176, 146] | 17.7 [-45.8, 117] |
| SS | -101 [-239, -35.3] | -25.3 [-38.6, -14.1] | 2.86 [-1.29, 5.86] |
| SM | -93.0 [-269, -27.7] | -12.6 [-23.4, -4.71] | -1.29 [-11.6, 4.71] |
| MS | -87.0 [-126, -21.4] | -18.8 [-43.2, 17.0] | -13.8 [-71.4, 7.0] |
| MM | -80.5 [-112, -58.3] | -19.3 [-31.8, -4.83] | -9.0 [-21.2, -1.83] |

**c-Fos expression**
| Group | Average number of c-Fos+ cells | Unpaired mean difference 'group' minus nsSS |
| --- | --- | --- |
| CC | 0.53±0.8 | -16.1 [-33.8, -9.39] |
| nsSS | 16.7±14.5 | 0 |
| SS | 25.6±11.6 | 8.97 [-8.99, 19.5] |
| SM | 13.1±11.0 | -3.56 [-21.7, 6.92] |
| MS | 17.8±15.7 | 1.10 [-13.1, 19.4] |
| MM | 37.9±29.7 | 21.2 [-3.53, 44.0] |

Overall mean of trained groups (nsSS and CC excluded) = 23.59±10.8

### 3.3. Experiment 3 - Effect of Robot avoidance and extinction on c-Fos protein expression

#### 3.3.1. Behavior

##### 3.3.1.1. Robot Avoidance Learning

Rats in aversively motivated groups SS, SM, MS, and MM learned to avoid the robot ((Fig. 15). Two-way ANOVA conducted on the number of shocks during training (test day excluded) revealed significant effect of Days (p< 0.0001, F(_3, 84_)=36.49), Group (p< 0.0001, F(_4, 28_)=21.05), and an interaction (p=0.0420, F(_12, 84_)=1.929). Holm-Šidák’s test showed that each experimental group received, among other, significantly less shocks on the last day of training compared to the first training day (SS and SM p<0.05; MM and MS p<0.005). No experimental group showed a statistically significant change between the third and the last training day. The active control nsSS group showed a high number of virtual shocks throughout its pseudo-training with no statistically significant difference between any of the sessions.

**Figure 15.**
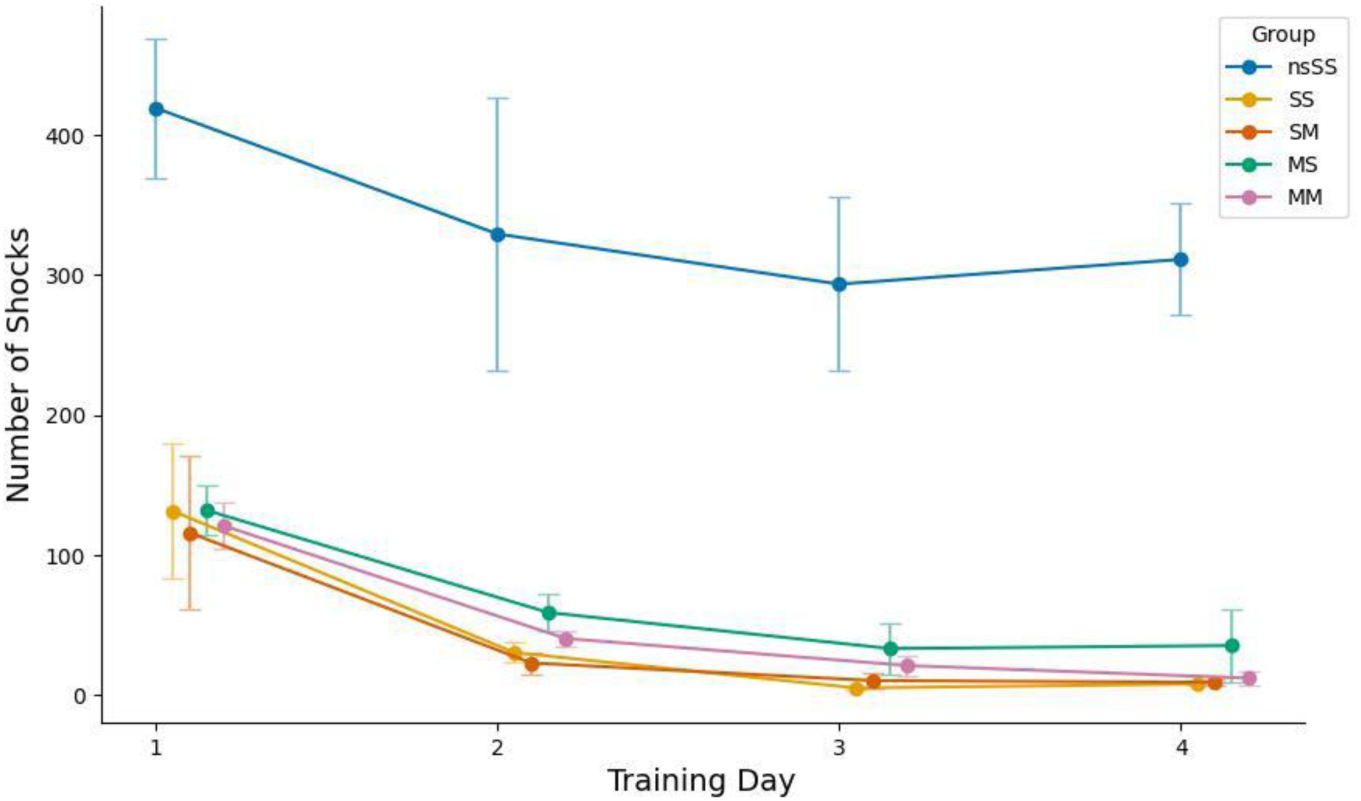
Robot Avoidance Training in Experiment 3. Training session lasted 20 min a day for four days. All experimental rats gradually reduced the number of shocks they received during the training period, whereas the no-shock active controls (nsSS) approached the robot substantially more (virtual shocks reported). Day1 > Day4 (p=0.0420). The overall mean for all experimental groups (nsSS excluded) on day 1 was 125.1±18.8 SEM, while on day 4 it was 16.2±6.9 SEM), Error bars are SEM.

##### 3.3.1.2. c-Fos expression

One-way ANOVA found an effect of groups on the mean number of c-Fos positive neurons (F_5,32_=21.51); p<0.0001. Tukey’s post-hoc test showed the CC group had a lower mean number of positive neurons than all other groups (p<0.0001), but no statistically significant differences were observed between the rest of the groups (Fig. 16). Representative images of c-Fos staining are in Fig. 17.

**Figure 16.**
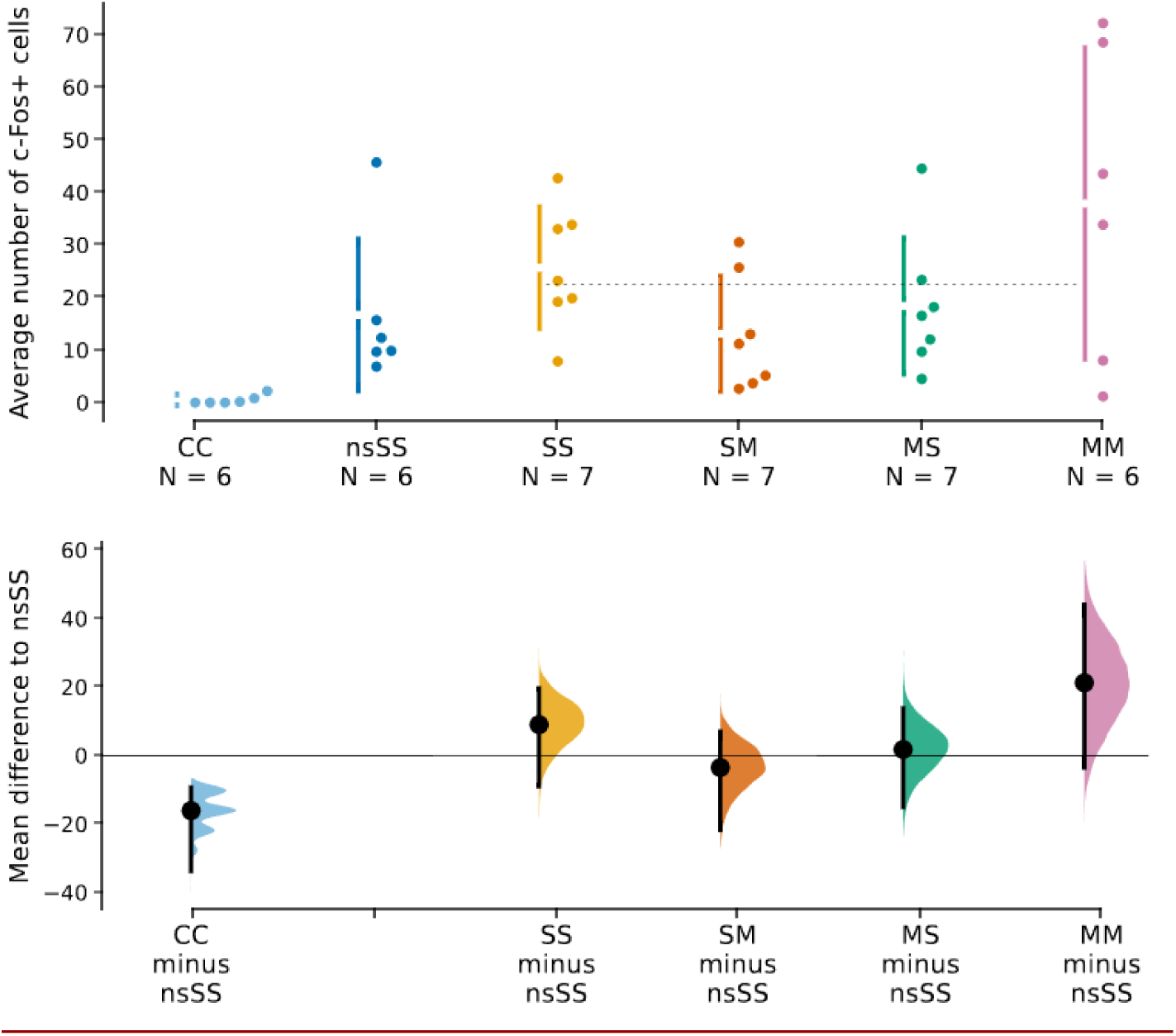
Average number of cells positive for c-Fos protein during the test session. **Top**: Raw data, the plot shows the average number of c-Fos+ cells for each animal, error bars are standard deviations with the gap representing the mean. The number of c-Fos+ cells in all trained (SS, SM, MS, MM) groups was significantly higher than in passive cage control CC animals (p<0.0001); no statistically significant differences were observed between the rest of the groups. The overall mean of trained groups (CC and nsSS excluded) was 23.59±10.8 std. The average number of c-Fos+ cells in the CC group was very low at 0.53±0.8. ‘Active groups’ > CC (p<0.0001); no significant differences between active groups. **Bottom:** Comparison of each group the nsSS group. Dots represent mean differences from the shared control of the CC group; vertical error bars are 95% confidence intervals with bootstrapped sampling distribution.

**Figure 17.**
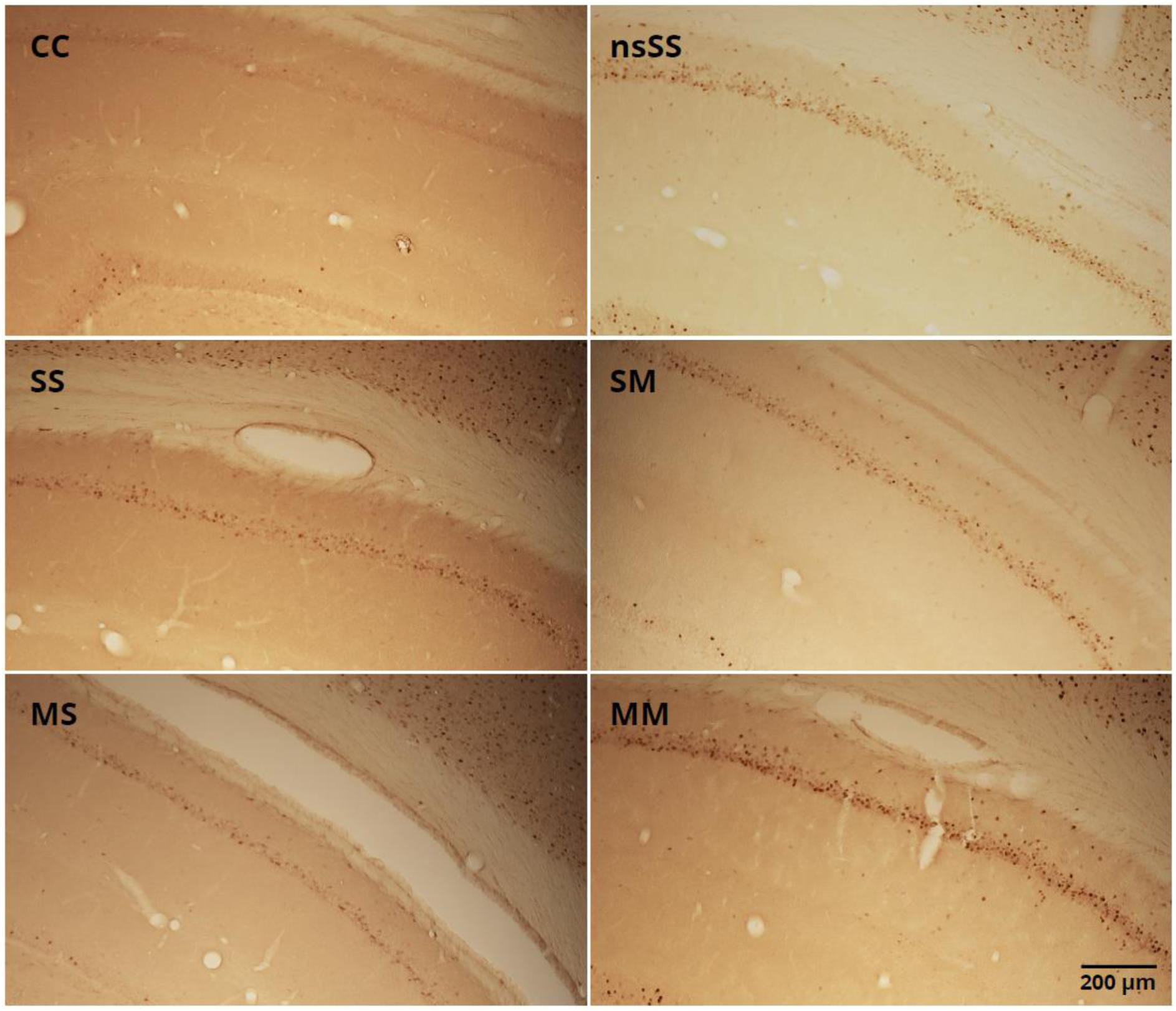
Representative images of c-Fos staining. c-Fos marks neural activity over the 30-minute test session. Few to none c-Fos+ cells present in dorsal CA1 of home caged controls (CC, left upper panel), while the nsSS, SS, SM, MS and MM groups showed similar amounts of neural activation.

#### 3.3.2. Behavioral Extinction and c-Fos expression

The paired mean difference of actual and virtual shocks between the last training day and the test day at 5 minutes (day 5 minus day 4, 95% CI) for each experimental group was: SS 9.29 [-1.57, 36.4], SM 19.6 [10.0, 43.4], MS -1.2 [-7.0, 2.8], MM 3.5 [-5.0, 19.0], nsSS 41.5 [-21.3, 193] (**Suppl. 5**).

The paired mean difference of actual and virtual shocks between the last training day and the test day at 20 minutes (day 5 minus day 4, 95% CI) for each experimental group was: SS 142 [50.1, 245], SM 89.9 [56.4, 177], MS 95.4 [-11.8, 237], MM 78.5 [32.8, 141], nsSS 113 [-4.67, 314] (Fig. 18).

**Figure 18.**
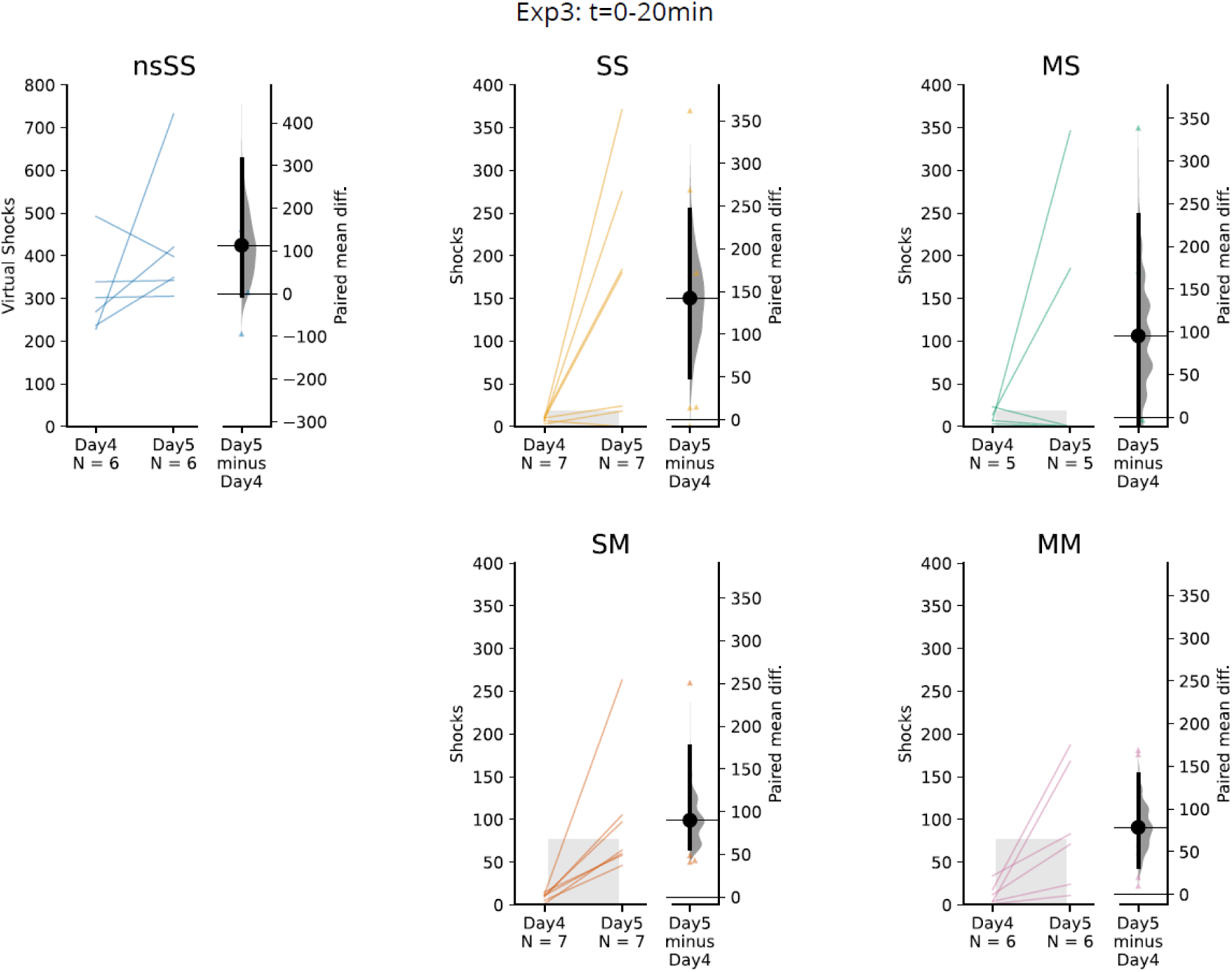
Actual and virtual shocks delivered in the first 20 minutes of the training day 4 and test day 5 and evaluation of behavioral extinction. Raw paired data of individual rats. Note the different y-axis on the left-hand side of the nsSS group. The right portion of each plot shows 95% CI for paired mean difference of day 4 and day 5 with distribution estimate. Grey bars represent one-sided tolerance intervals (TI, 99% proportion of population, 95% confidence level) later used as thresholds as described in the Methods section 2.2.5. The S-20 TI threshold is applied to groups with the robot stationary during the test (SS, MS); TI threshold M-20 is applied to groups with the robot moving during the test (SM, MM).

The TIs used for thresholds were the same as in Exp2. The resultant number of animals in each group was: Av-Av N=3; Av-Ext N=10; Ext-Ext N=3; Unclear N=9 (Fig. 19). The means and standard deviations for the average number of c-Fos+ cells (Fig. 20) were: nsSS 16.7±14.5; Av-Av 11.9±7.3; Av-Ext 17.6±16.9; Ext-Ext 35.9±6.8. In line with trends observed in Exp 2, the Ext-Ext group displaying the most of new extinction learning also displayed the highest c-Fos counts in accord with the notion that extinction learning stimulated plasticity-related IEG expression to accommodate new information and delayed the decline in IEG+ ensemble size observed after prolonged exploration in Exp1.

**Figure 19.**
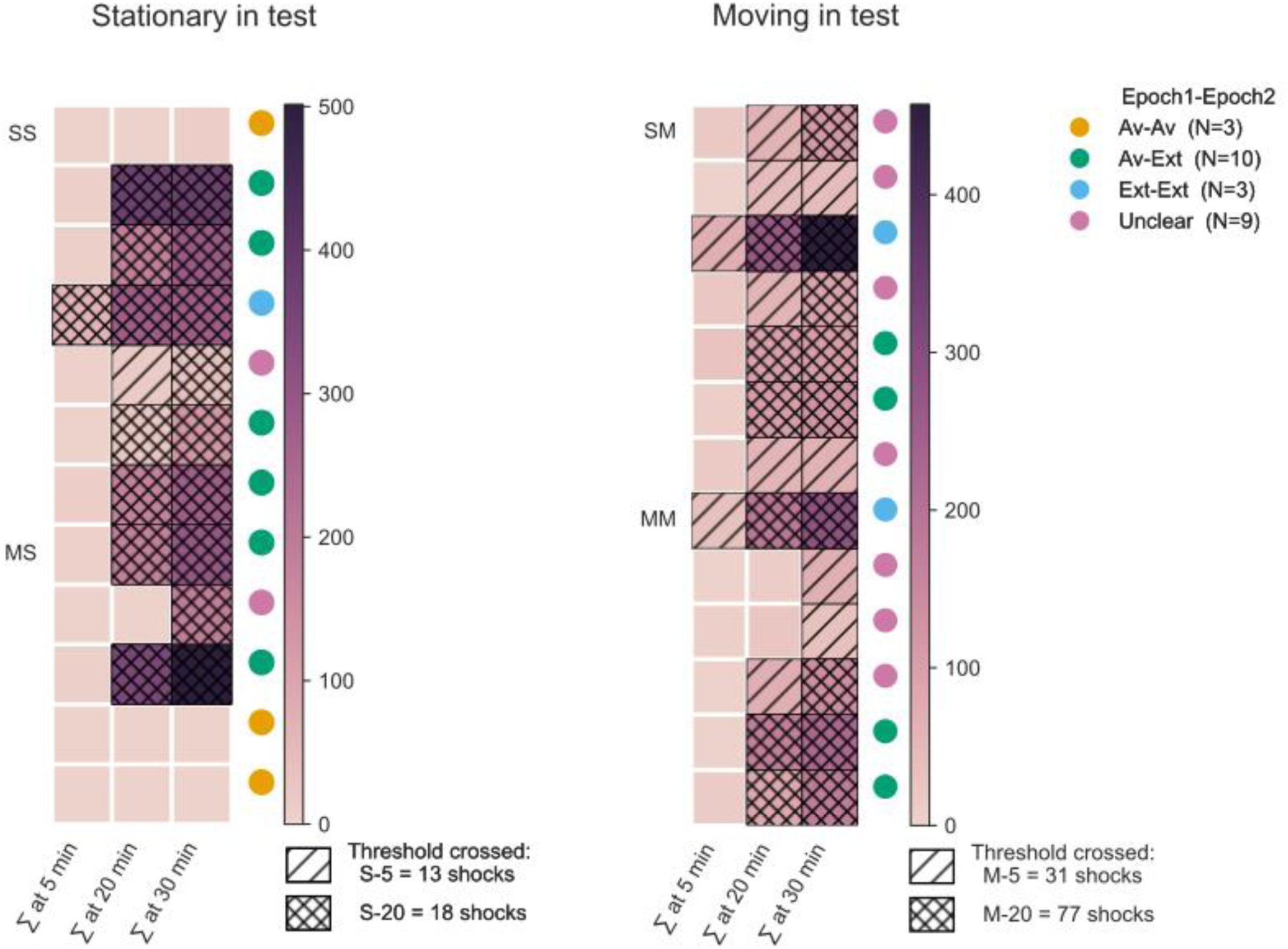
Avoidance and extinction groups in Experiment 3. Thresholds used to classify animals as ‘Av’ (avoidance) or ‘Ext’ (extinction) were the same as in Exp2. The ‘S’ and ‘M’ thresholds applied to groups with robot stationary (SS, MS), or moving during the test (SM, MM), respectively. The heatmap displays cumulative virtual shocks for each animal (y-axis) at given time points of the test (x-axis). Color bars represent the number of virtual shocks during the test. Hatchings mark a given threshold was exceeded at a given time point. Colored circles represent animals based on the presence of avoidance or extinction in Epochs 1 (first 5 min) and 2 (last 5 min).

**Figure 20.**
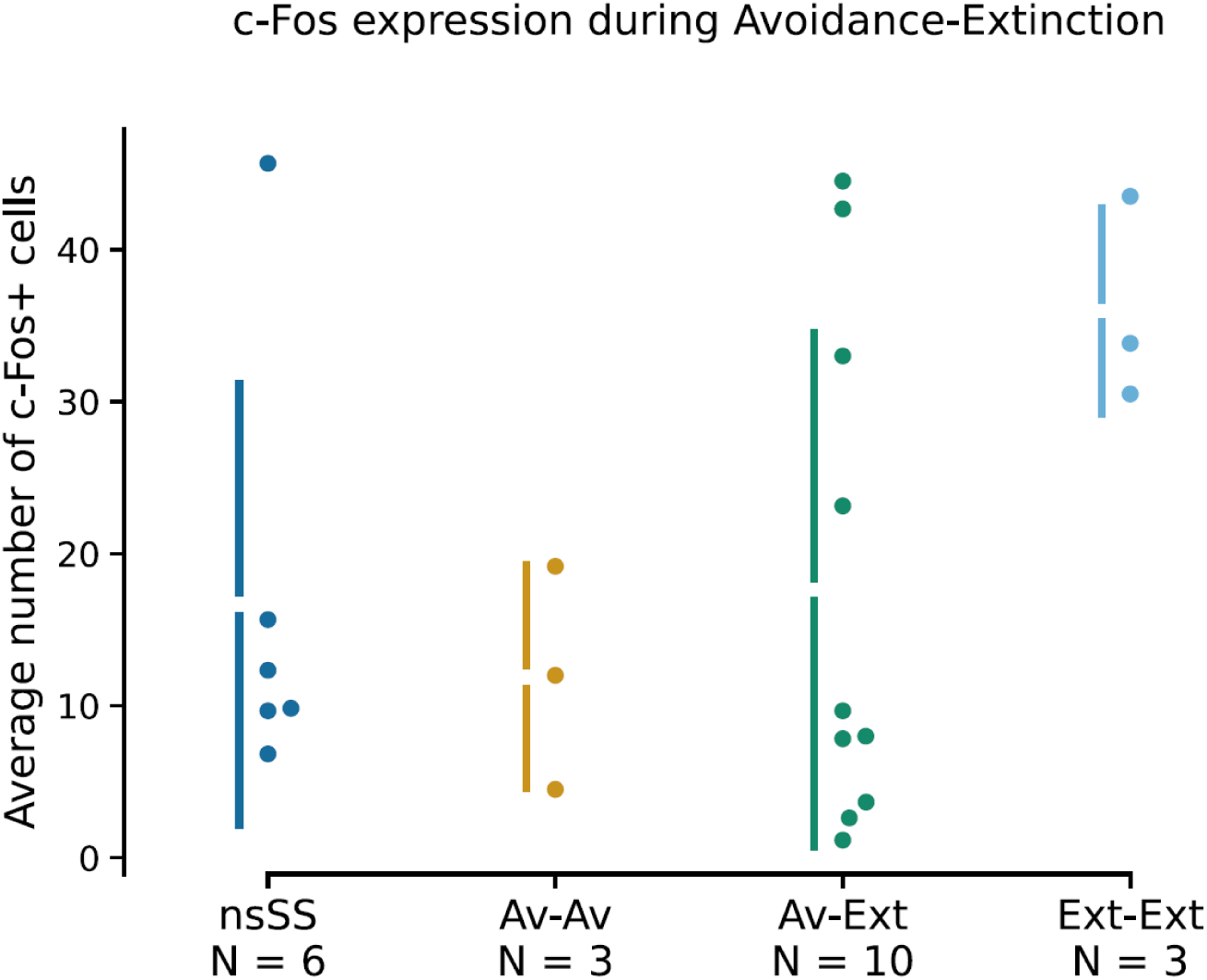
Average number of c-Fos+ cells relative to avoidance and extinction. Error bars are standard deviations, gaps represent means. Data markers represent individual animals. The Ext-Ext group displayed a notably higher number of c-Fos+ cells relative to the other groups, particularly the Av-Av.

## 4. DISCUSSION

### 4.1. A change of environment triggers context-specific response of IEG-expressing CA1 ensembles that declines over time

Exp1 showed that entering either a familiar or novel environment (context switch) triggers a wave of context-specific IEG expression, which fades over time. Also, our results corroborate the observations obtained using Ca^2+^ imaging (31) that CA1 ensembles activated closer in time are more similar to each other compared to ensembles activated further apart. We replicated the observation of contextual specificity of IEG+ ensembles in CA1 (ensemble similarity in groups A/A > A/B). The ensemble size and similarity in the A25/B group were practically identical to the A/B group, because the timing of their transfer to context B was exactly the same, resulting in a large *Arc+* ensemble with low similarity to the *Homer1a+* ensemble activated in context A at the start of the test. In contrast, the context switch occurred early for the A/B25 group, resulting in *Homer1a* expression in neurons activated by the early transfer to context B, some of which still express *Arc* by the end of the test, increasing the ensemble similarity. In fact, ensemble similarity in the A/B25 group was comparable to that of A/A, but the *Arc+* population was much smaller after the 25 min spent in context B. Similarly, only a fraction of neurons activated by entering context A at the start of the test also express Arc by its end in the A30 group. Our results show that the size of IEG+ ensembles declines if the animal persists in the environment for >25 minutes. The size of *Arc+* ensembles active during the last 5 minutes (Epoch2) was very low in animals that were already present in the environment (groups A/B25, A30); in fact, their *Arc+* ensemble size did not differ from the cage controls (CC). Overall, the IEG+ ensemble size reflects the recency of context switch regardless of familiarity with the context. On a longer timescale, the potency of entering a context to induce IEG expression may also decline, if many trials are massed on a single day (32).

Exp2 replicated findings of Exp1 that introduction to an environment (context switch) triggers a substantial IEG-ensemble response, which declines when animals remain in that context. This response did not diminish despite the rats being familiarized with the environment over four days of prior training. The ensemble sizes upon introduction to the novel and familiar contexts in Exps 1 and 2 were practically identical at ∼0.34 and ∼0.35 (average proportions of *Homer1a*+ neurons, Epoch1), respectively. This observation is in line with previous reports that spaced trials do not suppress the IEG ensemble response (32).

The decline in IEG ensemble size in Exp2 by ∼40% to ∼0.21 (average proportion of *Arc*+ neurons, Epoch2) was equivalent across all active groups, irrespective of the shock and robot motion during the test or training. The behavioral design of Exp2 was closest to the A30 group in Exp1 in that rats remained in one environment for the entire 30 minutes test. Although the ensemble size in Exp2 indeed declined compared to the Exp1 A/A, A/B, and A25/B groups (all ∼0.35 *Arc*+, Epoch2, no decline), the Exp2 ensembles were not quite as small as in groups A30 and A/B25 (∼0.12 *Arc*+, decline of ∼65%) of Exp1. The persistence of larger ensembles in Exp2 animals, despite the continuous presence in the same environment, might be attributed to the larger environment used in Exp2 compared to Exp1. The proportion of IEG-expressing neurons is greatly reduced in confined environments (33). The smaller environment used in Exp1 could have resulted in steeper ensemble size decline. The ensemble similarity in Exp2 was high (average similarity score 0.42) and almost identical in all groups. It was closer to that of groups A/A and A/B25 (∼0.44) rather than groups A/B (0.27) or A25/B (0.26) from Exp1.

### 4.2. Effect of task familiarity and novelty on IEG+ ensembles

Our previous study showed that the dorsal hippocampus is necessary for successful avoidance of moving, but not stationary, robot (23). We expected that increased hippocampal demand in groups with moving robot would lead to larger or more persistent ensemble sizes compared to groups with stationary robot. However, only minute differences in ensemble sizes of trained groups were observed in Epoch1: *Homer1a+* ensemble size ranged from 0.32 in SS to 0.37 in MM, and ∼0.35 in the incongruent SM and MS groups; practically no differences were present in Epoch2: *Arc+* was ∼0.18 in SS and ∼0.23 in the other experimental groups; and neither the ensemble similarity nor the c-Fos protein revealed any differences between the active groups. Previous research has shown that the amounts of IEG RNA and protein correlated with task performance (12,20,21), but this might be caused by variation in the IEG product content rather than in the number of expressing neurons, in accord with a previous report (22) that the proportion of dentate gyrus granule cells expressing Arc protein after active spatial avoidance task or mere exploration were equivalent.

Despite the lack of prominent differences between the active groups, all trained animals consistently displayed slightly smaller *Homer1a+* ensemble sizes (on average 0.35) in Epoch1) than the untrained nsSS group (0.39) – a relative difference of 12.5%. Although not statistically significant, this observation agrees with previous reports that IEG expression is lower during overtrained performance compared to initial acquisition [34–37]. However, no differences in *Arc+* ensemble size between trained and untrained animals were observed in Epoch2.

When we evaluated the IEG expression relative to avoidance/extinction in each epoch, we observed that animals avoiding the robot in Epoch1 (Av-Av, Av-Ext) displayed somewhat smaller *Homer1a+* ensembles (mean 0.31) compared to both the untrained nsSS group (0.39) and animals (Ext-Ext), in which extinction was present already in Epoch1 (0.38). This relative difference of ∼18% was larger than any observed between the original (robot training) groups, which did not discriminate avoidance from extinction. These observations suggest smaller ensembles are sufficient when performance becomes skilled, and that new extinction learning contributes positively to the ensemble size. This account is in accord with reported decreased IEG expression levels in overtrained animals (21) and increased IEG levels during behavioral extinction since extinction constitutes new learning (34).

The relative change in ensemble size between Epoch1 and 2 was relatively similar in the nsSS (-45%), Av-Av (-46%), and Av-Ext (-42%) groups. Notably, the smallest reduction between the two epochs was present in the Ext-Ext group (-31%). The smaller relative decrease in the Ext-Ext group might be attributed to more advanced learning in rats that discovered the absence of shocks early during the test. However, no prominent differences in ensemble overlap were observed between the groups.

### 4.3. IEG+ ensembles might be coupled to regularity between contextual change and uncertainty via novelty to promote timely memory formation

Why is the IEG ensemble dynamics characterized by sharp onset triggered by change of context followed by a decline? Why is this dynamic also true when revisiting a familiar environment? And why are hippocampal IEG+ ensembles contextually specific but mostly unaffected by (hippocampus-dependent) task demands? Since environments change over time, even re-entering a familiar context after a (sufficient) period of absence is a good indicator that updating may be required (4). Consequently, coupling IEG-expressing ensembles to contextual change would allow segmenting continuous experience into context-specific memory traces (35). The contextually specific IEG+ ensembles may act as indices (36) for individual memories (37–39) and context-relevant behaviors (5,40). The hippocampal index, by allowing reinstatement of cortical states, can aid in selection (41) of task-relevant behaviors, while itself remaining largely unspecific to task or behavioral details. We note that the view of neural systems coupled to law-like (probabilistic but regular) relation between contextual change and unexpected events is in line with the embodied and control loop approach to behavior and cognition (5,42–44), the elegant ‘subjective physics’ approach of Brette (45,46), and also consistent with the internally generated neural activity that provides initially meaningless frame of pattern that is later associated with experience (47–49). Moreover, this view seems plausible from an evolutionary perspective (50).

The coupling of IEG expression to contextual change in line with ‘behavioral tagging’ – an evolutionarily conserved phenomenon, where experience, which would otherwise only lead to a short-term memory, is consolidated in the long-term, if another, sufficiently novel experience causes proteosynthesis (16,51). Behavioral tagging is mediated by cellular and synaptic tagging and capture that involves IEG expression (51–53). **Novelty** can be not only absolute but also contextual (54–56), so that mere change of context might provide the novelty needed. The context change increases novelty and uncertainty; ensuing exploration and familiarization eventually reduce novelty as well as uncertainty. This pattern is matched by the recruitment of IEG-expressing ensembles. The dynamic range of *Arc* expression allows adaptive updating and enabling cognitive flexibility, whereas persistent expression leads to signal saturation, deficits in updating, and, consequently, inflexible behavior (2). The situation is akin to adaptation in the sensory system (57,58). When as many as nine sessions are massed on a single day, the novelty associated with a context switch fades and the IEG+ ensemble response is diminished (32).

## 5. CONCLUSIONS

As animals and neural systems evolved embedded in unstable environments, they exploited environmental regularities to extract knowledge and support adaptive behavior. Context aids prediction by narrowing down the range of options, despite the details of the forthcoming events remaining undetermined by context alone. Context change raises the expectations and triggers IEG+ ensemble recruitment, while task novelty modulates the endurance of contextually specific ensembles. The adaptive coupling of CA1 IEG+ ensembles to contextual novelty contributes to solving the stability-plasticity dilemma faced by animals living in a changing world.

## Author contribution: CRediT

Branislav Krajcovic: Conceptualization, Formal Analysis, Investigation, Writing – Original Draft, Writing – review & editing, Software, Visualization, Data curation, Methodology, Project administration

Daniela Cernotova: Investigation, Writing – Original Draft, Visualization, Project Administration

Ales Stuchlik: Methodology, Resources

Stepan Kubik: Methodology, Conceptualization, Writing – Original Draft, Writing – review & editing, Resources, Project administration, Supervision

Jan Svoboda: Methodology, Funding acquisition, Resources, Project administration, Data curation, Formal Analysis, Writing – review & editing, Supervision

## Acknowledgement

We would like to thank Hana Brozka for valuable discussions, Pavel Zitta for assistance with behavioral experiments, and Carl Olson for initial technical support with Python. The work was supported by IPHYS BIF – MEYS CR (Large RI Project LM2023050 Czech-BioImaging) and ERDF (Project No. CZ.02.1.01/0.0/0.0/18_046/0016045). Graphical abstract was created with BioRender.com. As non-native speakers we used ChatGPT to edit original text for brevity and readability, the authors reviewed and edited the content as needed and take full responsibility for the content of the publication.

## Research data

The raw data not present in the article will be made available by the authors upon request, without undue reservation.

## Funding

The work was supported by bilateral INTER-ACTION project LTAUSA19135 awarded to JS, and by the Czech Science Foundation grant GACR 21-16667K awarded to AS.

## Declaration of interest

Authors have no conflict of interest to declare.

## Notes

### Competing Interest Statement

The authors have declared no competing interest.

