## Supplements 1-5 for "CA1 ensemble plasticity is coupled to context change and modulated by task familiarity"

### Supplement 1:

#### Arc/Homer1a catFISH tissue processing and labeling

Immediately after the test session, rats were deeply anesthetized with isoflurane, brains were quickly (<2 min for Exp1, 2.5 min for Exp2) removed and flash-frozen in an isopentane bath equilibrated in dry ice/ethanol slurry. Rapid removal and freezing minimize confounding background expression induced by after-session stimuli or spontaneous activity. The brains were stored at -80°C and later cleaved into 4 mm segments using a tissue matrix. The segment containing the dorsal hippocampus was split by hemispheres, and blocks of 8 left hemispheres containing the dorsal CA1 were embedded in optimal cutting temperature medium (O.C.T.; Sakura, USA) and prepared for cryosectioning. The hemispheres were arranged in each block and sectioned together to maximize within-block and between-group comparisons. Twenty µm thick sections were cut in a cryostat (Leica CM1900 or Leica CM1850; Leica, Germany) at -17°C, mounted on gelatine-coated slides (Superfrost, Fisher), air dried, and stored at -80°C. All sample slides were processed simultaneously for each experiment with the double label (2 RNA probes) fluorescence *in situ* hybridization and DAPI counterstain as described previously (14,19,25). Briefly, both antisense riboprobes for *Homer1a* 3'UTR labeled by fluorescein and antisense, intron-enriched riboprobes for *Arc* labeled by digoxigenin were used for a single overnight hybridization and then sequentially detected by secondary antibodies: anti-fluorescein-HRP (Jackson labs) and anti-digoxigenin-HRP FAB fragments (Roche). The signals were then developed using tyramide amplification systems (TSA, Perkin Elmer, USA): TSA-Fluorescein for *Homer 1a* probes and TSA-Cy3 for *Arc* probes. Afterward, the slides were counterstained with DAPI (Invitrogen), coverslipped with antifade medium (Vectashield, Vector labs), and sealed with nail polish.

### Supplement 2:

#### *Arc*/*Homer1a* catFISH Image Acquisition and Sampling

All images were acquired on an inverted Leica TCS SP8 AOBS WLL confocal laser scanning microscope: 20×oil immersion objective; blue signal DAPI, 405nm excitation, 415-490nm bandpass; green signal TSA-Fluorescein, 488nm excitation, 510-550nm bandpass; orange/red signal TSA-Cy3, 561nm excitation, 610-680nm bandpass. Excitation settings (laser power, gain, and offset) were set the same for the whole slide but varied slightly between slides due to the slight variation in staining strength in each slide; excitation settings for a given slide were based on FOV with the strongest signal in each IEG channel so that approximately three IEG signal points (intranuclear foci) reaching saturation were present in this FOV - this protocol yields standardized and reproducible excitation settings, increasing the comparability between slides.

### Supplement 3:

#### Exp2 cell counts

For the experimental groups, 448 Z-stacks in total were analyzed, 32 Z-stacks were removed due to tissue or cell damage; in total, 17759 nuclei were analyzed for experimental groups, and an estimated 1268 nuclei or 6.7% were not included in the analysis (estimate is based on the average number of nuclei analyzed per Z-stack, making the estimated total of 19027 nuclei for all experimental groups pooled); the average number of cells analyzed per experimental group was 3552 (estimated 253 nuclei per experimental group were missing from the analysis); an average number of nuclei analyzed per experimental animal was 444 (estimated 31 nuclei per experimental animal were missing). For the test day cage control animals, 93 Z-stacks were analyzed, and 3 Z-stacks were removed due to tissue damage.

#### Supplement 4:

#### DAB c-Fos staining

Free-floating slices were washed in 1x PBS, then incubated for 30 min in peroxidase blocking serum (1x PBS with 10% MeOH and 3% $H_2O_2$), followed by 30 min incubation in blocking serum (1x PBS with 5% serum and 0.3% Triton). The slices were then incubated overnight on a shaker at 4 °C (1:1000 c-Fos antibody, #AB190289, Abcam, with 5% serum and 0.3% Triton). On the second day, slices were washed thoroughly in 1x PBS and incubated in biotinylated anti-rabbit secondary antibody (1:500, #SAB4600006, Sigma Aldrich) for 2 hours, then with avidin-biotin complex (9 µl of each / 1 ml PBS, #R37627, Invitrogen) for 1 hr. The DAB staining was finalized using 10 mg Dab (#D5905, Sigma Aldrich) dissolved in 50 ml PBS, and by adding 15µl 30% $H_2O_2$ right before staining. The final staining was stopped after 6–10 min, the slices were washed with tap water, then 1x PBS. The slices were mounted on gelatin-coated slides, soaked in Neo-Clear, and coverslipped using Neo-Mount.

### Supplement 5:

#### Comparison of actual and virtual shocks on day 4 and day 5 at t=0-5min in Experiment 3

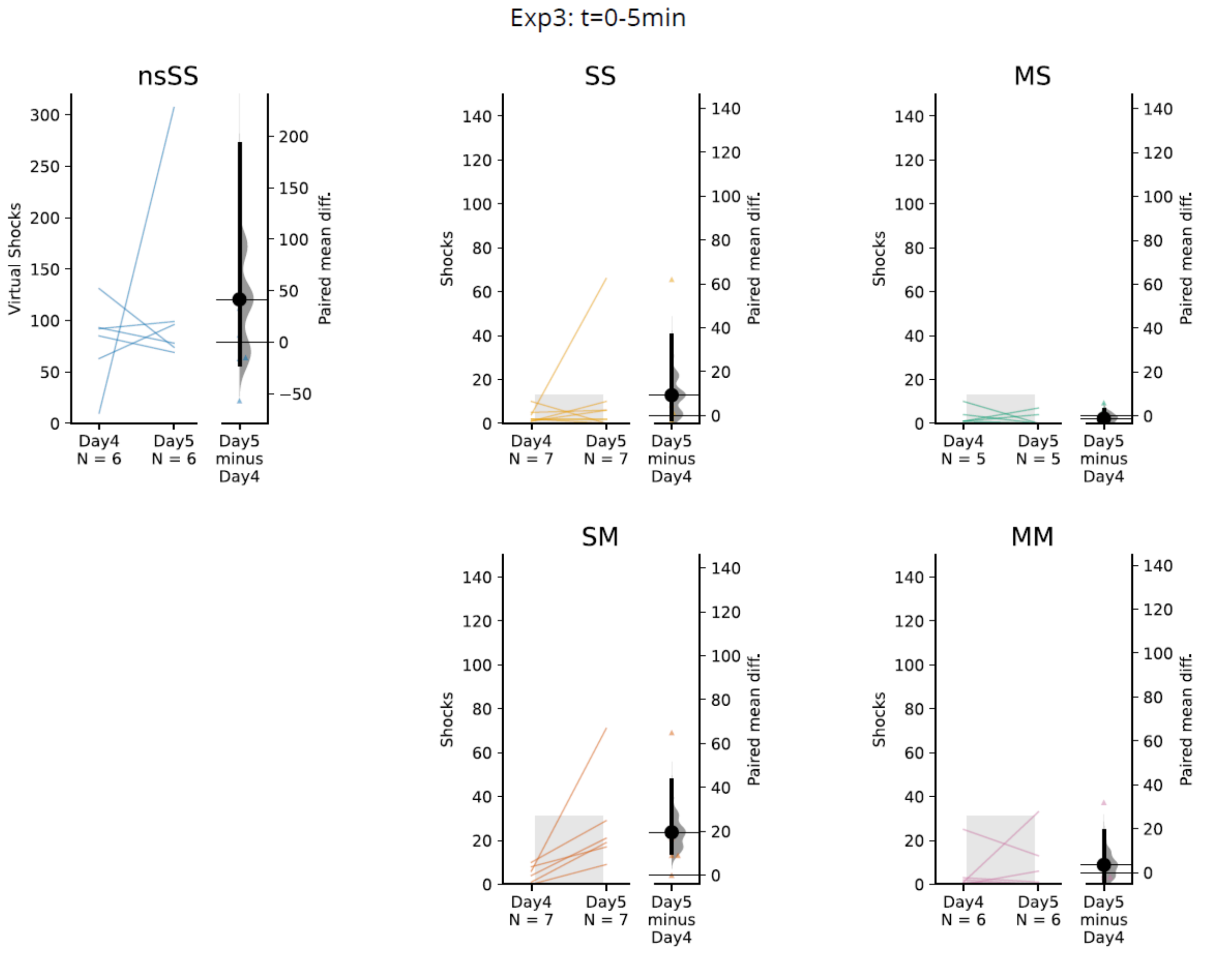


**Supplement 5 Figure: Actual and virtual shocks delivered in the first 5 minutes of the training day 4 and test day 5.** Raw paired data of individual rats depicting the number of actual and virtual shocks received during the first 5 minutes of training day 4 and test day 5, respectively. Note the different y-axis on the left-hand side in the nsSS group. The right portion of each plot shows 95% CI for paired mean differences of day 4 and day 5 for the whole group with distribution estimate. Gray bars represent one-sided tolerance intervals (TI, 99% proportion of population, 95% confidence level) later used as thresholds as described in the Methods. The S-5 TI threshold is applied to groups with robot stationary during the test (SS, MS); TI threshold M-5 is applied to groups with robot moving during the test (SM, MM).
